# Positron emission tomography (PET) tracer enables imaging of high CD73 expression in cancer

**DOI:** 10.1101/2024.11.29.625970

**Authors:** Clemens Dobelmann, Constanze C. Schmies, Georg Wilhelm Rolshoven, Mirko Scortichini, Stefan Wagner, Andreas Isaak, Riham M. Idris, Jennifer Dabel, Lucie Grey, Karolina Losenkova, Susanne Moschütz, Antje Keim, Sandra Höppner, Jouko Sandholm, Pia Boström, Maija Hollmén, Norbert Sträter, Sven Hermann, Gennady G. Yegutkin, Kenneth A. Jacobson, Sonja Schelhaas, Christa E. Müller, Anna Junker

## Abstract

Ecto-5’-nucleotidase (CD73) is a potential new drug target for cancer immunotherapy. Its overexpression is associated with various aggressive cancers, including triple-negative breast cancer (TNBC) and pancreatic cancer, making it a promising target for diagnostic imaging. Besides antibodies, small molecule CD73 inhibitors have been developed and are currently in clinical trials. This study aimed to develop and evaluate fluorine-18 labeled high-affinity CD73 inhibitors as tracers for the non-invasive positron emission tomography (PET) imaging of CD73 expression in cancer. Two CD73 inhibitors were selected for radiolabeling based on their high potency (K_i_ values of ca. 1 nM), and favorable pharmacokinetic properties providing [^18^F]PSB-19427 ([^18^F]**1**) and [^18^F]MRS-4648 ([^18^F]**2**). Ex vivo imaging studies on human breast cancer tissues indicated specific binding of both radiotracers. Subsequent in vivo studies proved [^18^F]**1** to be superior due to its long elimination half-life and its accumulation in TNBC and pancreatic cancer tissues, suggesting its potential as a versatile PET tracer for imaging various solid tumors. [^18^F]**1** significantly outperformed [^18^F]FDG in visualizing triple-negative breast cancer, offering potential advantages over [^18^F]FDG in terms of specificity and diagnostic accuracy. Thus, [^18^F]**1** is a PET tracer with outstanding properties suitable for broad application in cancer diagnosis and potentially in therapy control. Based on these results, further clinical development of PET tracers targeting CD73 is warranted.

**One Sentence Summary:** A PET tracer for imaging CD73 expression was developed, enabling cancer diagnosis.

## INTRODUCTION

Human ecto-5’-nucleotidase, also termed cluster of differentiation 73 (CD73), is a ubiquitously expressed homodimeric enzyme that exists in soluble and membrane-(glycosylphosphatidylinositol)-bound form. Additionally, it is commonly found on the surface of exosomes (*1*). CD73 catalyzes the extracellular hydrolysis of nucleoside monophosphates, yielding the corresponding nucleosides. CD73 converts extracellular AMP to immunosuppressive adenosine and inorganic phosphate(*2, 3*). AMP is generated from pro-inflammatory ATP or other adenine nucleotides(*4, 5*). Due to the hypoxic tumor microenvironment (TME), the gene NT5E encoding for CD73 is upregulated in cancer cells(*6*). Additionally, upregulated pro-inflammatory factors such as TGF-β, interferons (IFNs), TNF, IL-1β, and the Wnt/β-catenin pathway activation further promote CD73 expression(*7*). The enzyme is expressed on cytotoxic CD8 T cells (*1*) and regulatory T cells (Tregs)(*8*) and is overexpressed in various cancer types, including melanoma, bladder, colon, ovarian, pancreatic, and breast cancer, and was shown to promote cancer cell migration, invasion, epithelial-mesenchymal transition (EMT), and possibly contributes to chemotherapy resistance(*9–13*). Several studies have demonstrated the prognostic value of CD73 expression in triple-negative breast cancer (TNBC)(*14*), pancreatic cancer (PC)(*15*), and lung adenocarcinoma(*16*). Various clinical phase I/II trials with small molecule CD73 inhibitors (AB680/Quemliclustat, LY3475070), and anti-CD73 monoclonal antibodies (MEDI9447/Oleclumab, NZV930/SRF373, TJ004309/TJD5, CPI-006, BMS-986179) are ongoing(*17, 18*). In recent years, based on our lead structure PSB-12379(*19*), a variety of nucleotide-derived CD73 inhibitors with nanomolar potency at rodent and human CD73, high selectivity, and high metabolic stability have been developed and tested as potential anticancer drugs(*20–26*).

Given the general role of CD73 in cancer development and progression, we were interested in the diagnostic potential of positron emission tomography (PET) imaging of CD73 expression in cancer, particularly in TNBC and pancreatic cancer. Breast cancers show a very high incidence comprising ca. 12 % of all cancers(*27*). TNBC accounts for 10-20% of all breast cancers and exhibits an especially aggressive clinical progression(*28*). Early and precise detection of breast cancer is expected to enhance the odds of survival. Especially for TNBC, many standard imaging methods such as mammography and ultrasound fail since detectable abnormalities are missing, whereas CD73 is considered a promising biomarker for TNBC (*10, 29*).

Due to its asymptomatic progression at early stages and its tendency to form distant metastases early on, pancreatic cancer is particularly aggressive and among the deadliest cancers (5-year survival rate <9%)(*30*). Despite much effort, there is still no effective drug available for pancreatic cancer therapy, and the only potentially curative treatment to date is its complete surgical resection. Hence, the key to optimal therapy management is tumor detection at a very early stage. PET with 2-deoxy-2-[^18^F]fluoro-D-glucose ([^18^F]FDG) as the commonly employed radiotracer for the imaging of pancreatic cancer leads to 90 % overall diagnostic accuracy but with low spatial resolution. Moreover, false-positive signals caused by physiologic FDG uptake limit the detection of small metastases(*31*). New, reliable tumor markers are urgently needed for the successful diagnosis of pancreatic cancer. Most recently, encouraging preclinical and clinical results of phase I and II studies of CD73 inhibitor AB680 (Quemliclustat) in pancreatic cancer were reported (NCT03677973, NCT04575311 NCT04104672, NCT05915442). A phase II clinical study of Zimberelimab and Quemliclustat in combination with chemotherapy in patients with borderline resectable and locally advanced pancreatic adenocarcinoma (NCT05688215) is currently ongoing. This highlights the great potential of targeting CD73 in pancreatic cancer.

Thus, CD73 appears to be a promising biomarker for both TNBC and pancreatic cancer, worth of in-depth evaluation via PET imaging. In the present study, we developed two structurally diverse fluorinated CD73 inhibitors, the adenine-based PSB-19427 (**1**) and the cytosine-based MRS-4648 (**2**). They were evaluated for their CD73 potency and accumulation in biopsy tissue samples from breast cancer patients, demonstrating high CD73 affinity (Fig. 1, Scheme 1). Radiofluorination with fluorine-18 led to the corresponding PET tracers [^18^F]PSB-19427 ([^18^F]**1**) and [^18^F]MRS-4648 ([^18^F]**2**). [^18^F]**1** displayed promising biodistribution and metabolic stability in vivo. In xenograft mouse models, [^18^F] **1** significantly outperformed [^18^F]FDG in detecting TNBC tumors, providing a much higher tumor-to-background signal. Additionally, [^18^F] **1** effectively visualized pancreatic cancer tumors, suggesting its potential as a versatile PET tracer for imaging various solid tumors.

**Fig. 1.**
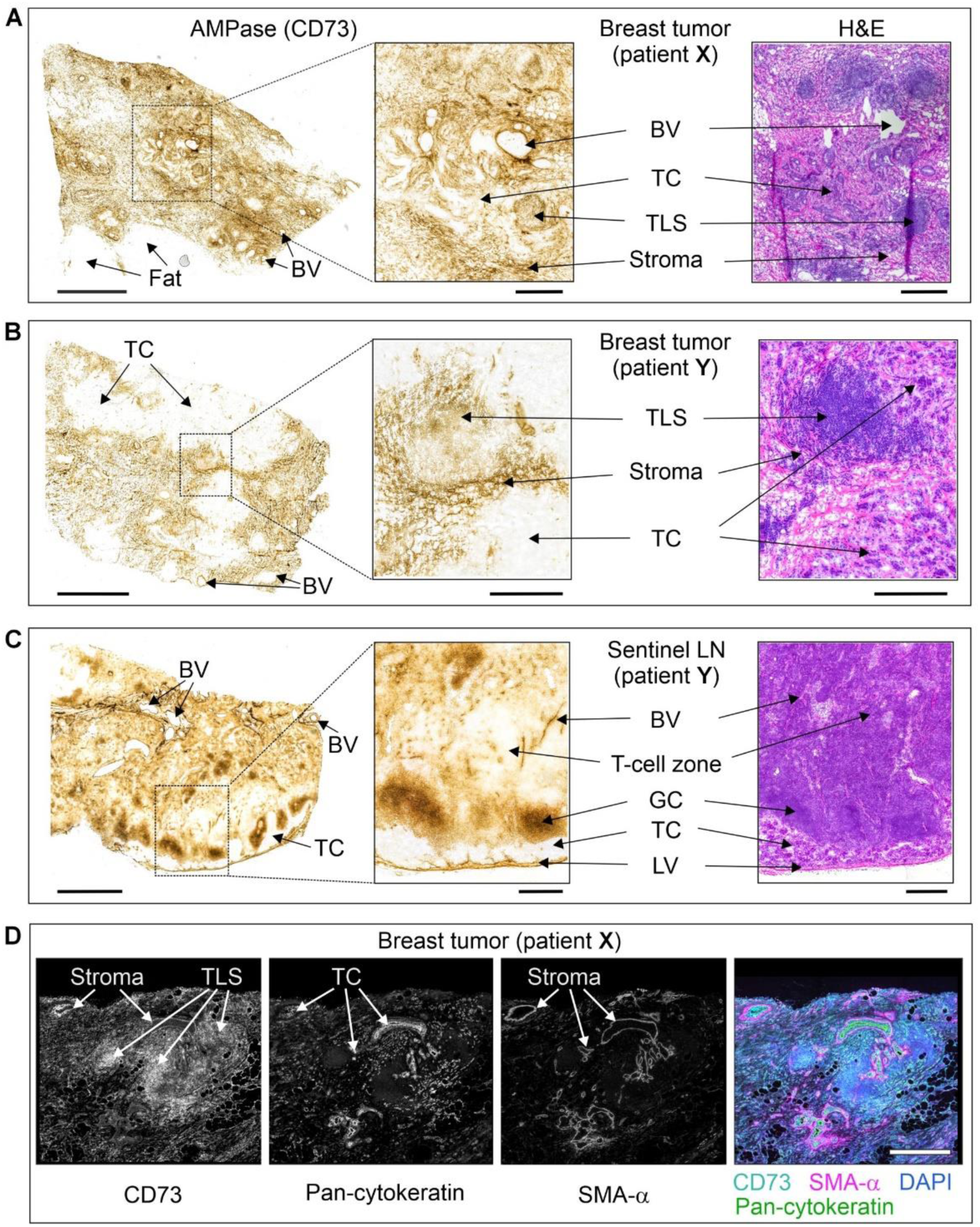
Tissue-specific distribution of CD73 in the breast tumor and lymph nodes. The primary tumors and sentinel lymph nodes (LN) were obtained from two patients with hormone-receptor-positive/Her2-negative grade II infiltrating ductal carcinoma (patient **X**), and hormone-receptor-negative/Her2-positive grade III infiltrating ductal carcinoma with micropapillary differentiation (patient **Y**). CD73 activity was assayed by incubating the cryosections of breast tumors from patients **X** (A) and **Y** (B), and also sentinel LN from patient **Y** (C) with 400 µM AMP in the presence of Pb(NO_3_)_2_, followed by microscopic detection of AMP-derived inorganic phosphate (P_i_) as a brown precipitate of formed Pb_3_(PO_4_)_2_. Tissue sections were also stained with hematoxylin and eosin (H&E). (D) For immunofluorescence staining, the tumor section from patient **X** was co-stained with an anti-CD73 antibody, together with a marker of epithelial cancer cells (pan-cytokeratin) and the stromal marker α-smooth muscle actin (SMA-α). Single channels are shown in grayscale, and the right panel displays a merged image with nuclei counterstained with DAPI. Abbreviations: BV, blood vessels; GC, germinal center; LV, lymphatic vessels; TC, tumor cells, TLS, tertiary lymphoid structure. Scale bars, 2 mm (A-C) and 500 µm (A-C, right insets, D).

**Scheme 1:**
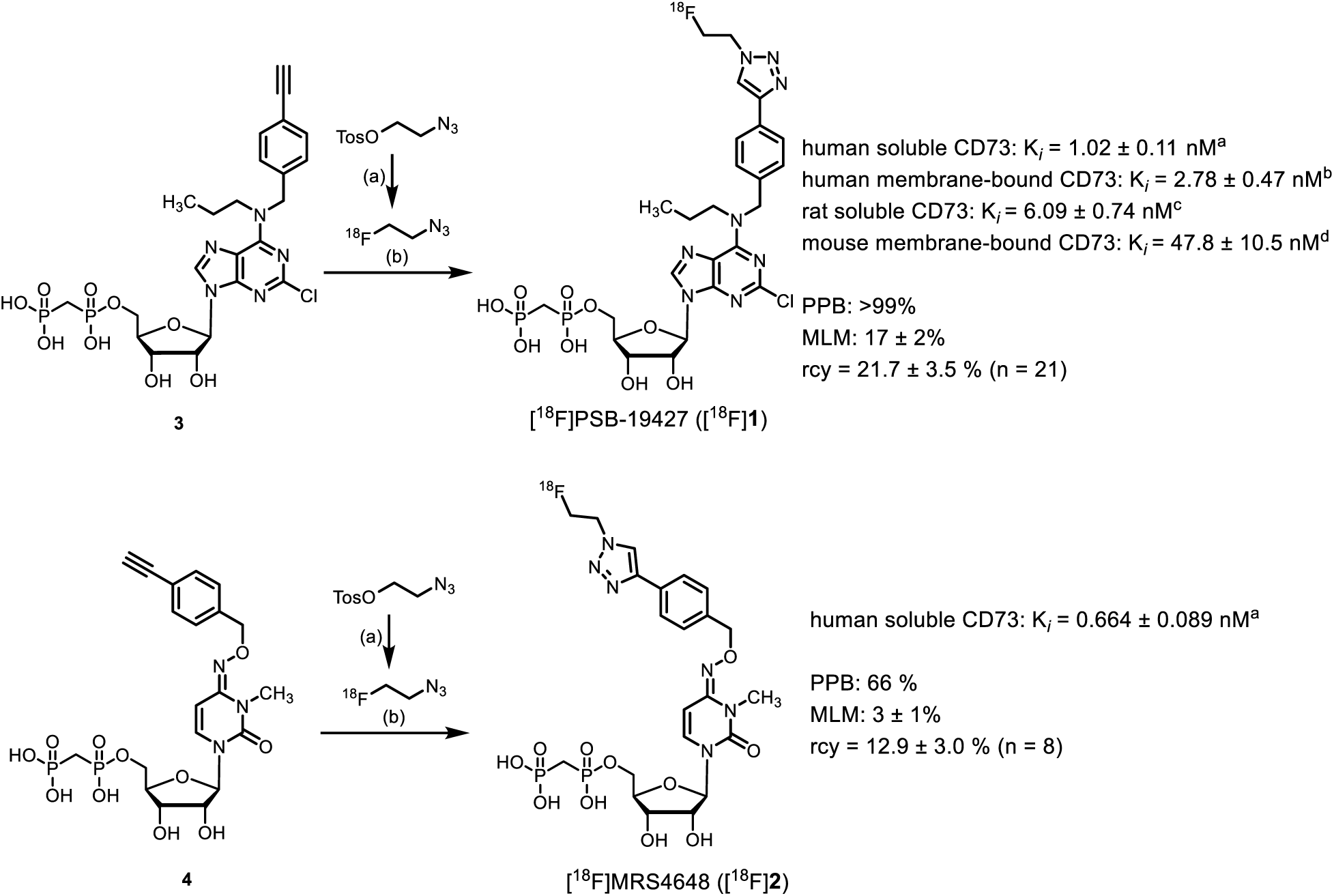
Radiosynthesis and in vitro assay data of [^18^F]PSB-19427 ([^18^F]1) and [^18^F]MRS-4648 ([^18^F]2). Reagents and conditions: (a) [^18^F]KF, K_2.2.2_, K_2_CO_3_, ACN, 110 °C, 3 min. (b) CuSO_4_, sodium ascorbate, DMF/HEPES buffer, 60 °C, 30 min. In vitro CD73 assays: ^a^Soluble recombinant human CD73 (K_m_17 µM, substrate AMP (5 µM)); ^b^Membrane preparation of human TNBC (MDA-MB-231 cell line, K_m_ 14.8 µM, substrate AMP (5 µM)); ^c^Soluble recombinant rat CD73 (K_m_ 59 µM, substrate AMP (5 µM)); ^d^Membrane preparation of mouse 4T1.2 breast cancer cell line (K_m_ 67.6 µM, substrate AMP (5 µM)). MLM, mouse liver microsomes; rcy, radiochemical yield.

## RESULTS

### Design and development of PET tracers

Several target compounds were designed based on our previous structure-activity relationship studies(*19, 24*), enabling the introduction of a fluorine atom in the final reaction step. Two structurally diverse inhibitors, PSB-19427 (**1**, K*_i_* = 1.02 ± 0.11 nM, human CD73) and MRS-4648 (**2**, K*_i_* = 0.664 ± 0.089 nM, human CD73) (for synthesis, see Supporting Information, Schemes S1 and S2), were characterized as highly potent competitive inhibitors of human CD73 with K*_i_* values of around 1 nM, determined using recombinant soluble human CD73 in an enzyme inhibition assay(*32*). Furthermore, for PSB-19427 the CD73 affinity at rat CD73 (K*_i_* = 6.09 ± 0.74 nM, rat soluble CD73) and mouse CD73 (K*_i_* = 47.8 ± 10.5 nM, mouse membrane-bound CD73) was determined. Both compounds were additionally evaluated in vitro for their metabolic stability in mouse liver microsomes (MLM) and plasma protein binding (PPB). PSB-19427 (**1**) displayed higher PPB (>99% but lower microsomal stability (17% decomposition in MLM after 90 min incubation in comparison to MRS-4648 (**2**, 66% PPB, 3% decomposition).

### PSB-19427 (1) and MRS-4648 (2) bind to CD73 in breast cancer tissues with high affinity and inhibit enzymatic activity

The binding of the non-radioactive CD73 inhibitors was further evaluated in primary breast tumor tissues and sentinel lymph nodes (LN) surgically removed from two patients with hormone-receptor-positive/Her2-negative grade II infiltrating ductal carcinoma (patient **X**) and hormone-receptor-negative/Her2-positive grade III infiltrating ductal carcinoma with micropapillary differentiation (patient **Y**). Tissue-specific distribution of AMPase activity (hydrolysis of AMP to adenosine and phosphate) and expression levels of CD73 in the breast tumor and the TME were determined by lead nitrate-based enzyme histochemistry, where the lead phosphate that precipitated due to CD73 activity was visualized as a brown deposit (Fig 1A-C), and by immunofluorescence staining (Fig. 1D), respectively (*33*). Additional staining of the tissue cryosections with haematoxylin and eosin enabled the visualization of the main histological structures (Fig. 1A-C, right insets)(*26*). Both AMPase activity and CD73 immunoreactivity were primarily associated with SMA-α^+^ stromal cells, tertiary lymphoid structures (TLS, mainly comprised of T- and B-cell aggregates(*34*)), blood and lymphatic vessels, and B-cell zone (in case of sentinel LN dissected from patient **Y**), but not with pan-cytokeratin-positive tumor cells themselves.

Subsequent competitive analysis of CD73-mediated AMPase activities was performed by incubating tissue cryosections with AMP and CD73 inhibitors at different concentrations. Fig. 2 depicts representative images of AMP-specific staining in breast tumors from patient **X** (panel A) and patient **Y** (panel B), determined in the absence (control) and in the presence of 10 nM of the potent CD73 inhibitor PSB-19427 (**1**). Treatment of tissue cryosections with increasing concentrations of PSB-19427 (**1**) and MRS-4648 (**2**) (10-50 nM), but not with the employed concentration of the classical, much less potent CD73 inhibitor AMPCP (**5**, 50 nM), reduced CD73/AMPase activity in the breast TME by ∼60-70% (Fig. 2C).

**Fig. 2.**
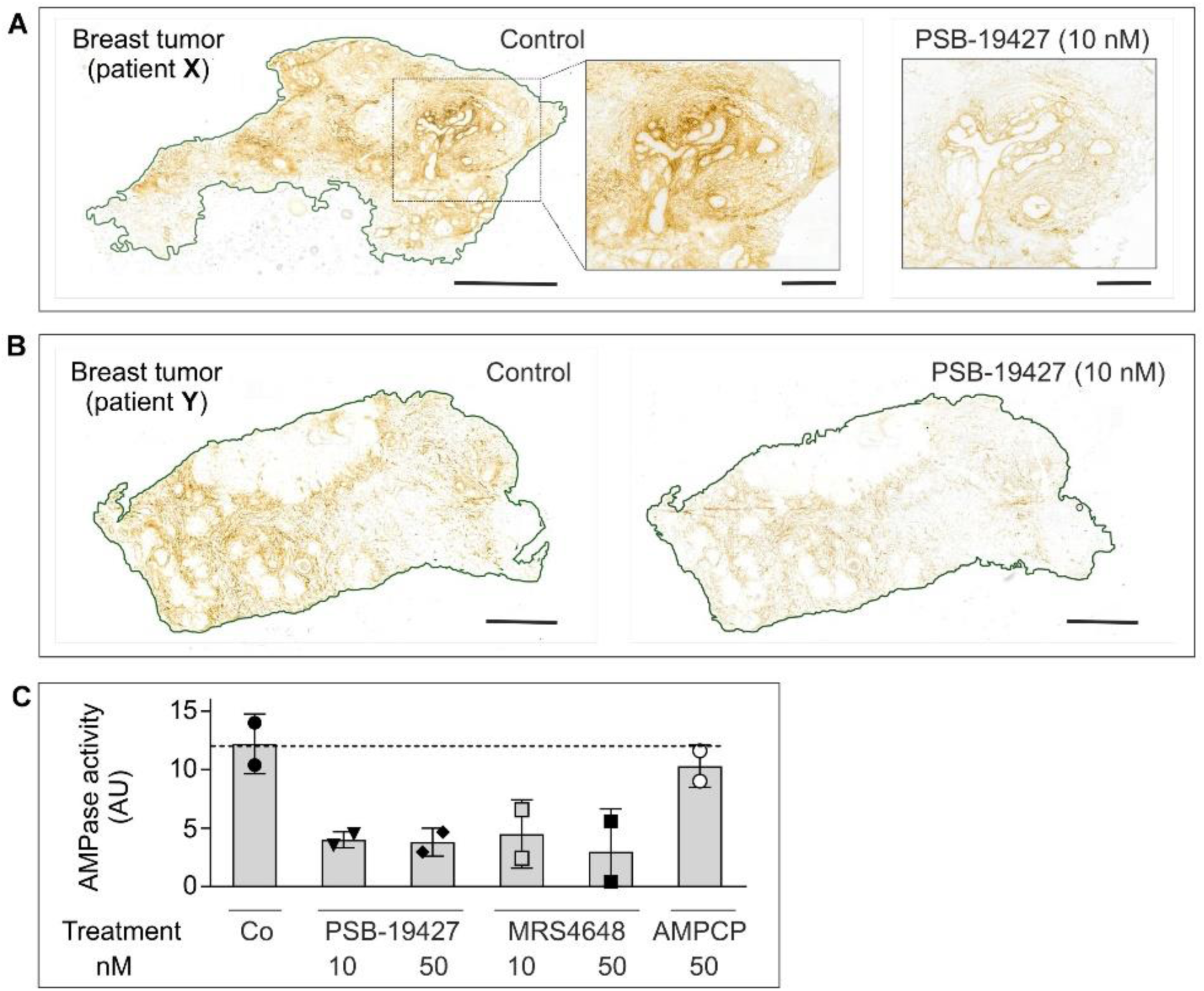
Effects of CD73 inhibitors on AMPase activity in human breast tumor tissues. The effects of CD73 inhibitors on AMPase activity was determined in situ by incubating primary breast tumor samples from patient X (A) and patient Y (B) with 400 μM AMP and 1.5 mM Pb(NO_3_)_2_ in the absence (control) and the presence of the indicated concentrations of CD73 inhibitors. (**C**) Mean pixel intensities of brown staining due to the hydrolysis of AMP leading to Pb_3_(PO_4_)_2_ precipitation were quantified in the selected regions and expressed as arbitrary units (AU) (mean ± SEM; n = 2). Scale bars: 2 mm (A, B), and 500 μm (A, right insets).

Based on these encouraging data, we decided to proceed with both compounds to radiosynthesis, autoradiography and in vivo PET imaging studies.

### Radiosynthesis

CD73 inhibitors **1** and **2** were subsequently prepared in ^18^F-labeled form. The alkyne-substituted precursors **3** and **4** were subjected to an azide-alkyne Huisgen-cycloaddition reaction using ^18^F-labeled fluoroethyl azide (Scheme 1). [^18^F]PSB-19427 ([^18^F]**1**) was obtained in 21.7 ± 3.5 % radiochemical yield (rcy), with >99% radiochemical purity (rcp) and a molar activity (A_m_) of 2.3-54.3 GBq/µmol (n = 21). [^18^F]MRS-4648 ([^18^F]**2**) was isolated in 12.9 ± 3.0 % rcy, with >98% rcp and a molar activity of 0.4-6.3 GBq/µmol (n = 8). Radiochemical identity was confirmed by observing comobility upon spiking with authentic PSB-19427 (**1**) and MRS-4648 (**2**) samples, respectively, in radio-high-performance chromatography. Both tracers displayed high affinity to glass surfaces but were sufficiently soluble in water for injection (WFI)/ethanol (9:1) at concentrations required for imaging applications (50-150 MBq/mL).

### Lipophilicity (log*D*_7.4_) and stability of radiotracers

The log*D*_7.4_-values of both radiotracers were determined by a previously described method based on their distribution between phosphate-buffered saline (PBS) and octan-1-ol(*35*). This revealed the higher polarity of the purine-based [^18^F]PSB-19427 ([^18^F]**1**; log*D*_7.4_ = -0.12 ± 0.03 as compared to the pyrimidine-derived [^18^F]MRS-4648 ([^18^F]**2,** log*D*_7.4_ = 0.74 ± 0.29). Both tracers were found to be stable for at least 90 min at room temperature in human and mouse plasma (100%, see Fig S1).

### Autoradiography of [^18^F]PSB-19427 and [^18^F]MRS-4648 in human breast cancer samples

To test the specific radiotracer binding to its target, we performed in situ autoradiography. [^18^F]MRS-4648 ([^18^F]**2**) was applied onto cryosections of neighboring slides of tumor tissues from primary breast tumors from patient **X** (Fig. 3A) and patient **Y** (Fig. 3B), used for immunohistochemistry and AMPase activity determination. Sentinel LN from patient **Y** was incubated with [^18^F]PSB-19427 ([^18^F]**1**) (Fig. 3C). In all three cases, a significant accumulation of radiotracer was detected in the breast tumor tissue (Fig. 3A-C, left panels). Notably, residual activity was also detected outside the tissue boundaries, which presumably reflects the ability of the radiotracers (especially [^18^F]**2**) to bind to the microscope slide. Importantly, co-incubation of the samples with a ∼1000-fold excess of the standard CD73 inhibitor AMPCP (**5**), or our recently developed pyrimidine-based CD73 inhibitor JMS0414 (**6,** Fig. 3D)(*24*) with the radiotracer markedly reduced the amount of tissue-associated radioactivity (Fig. 3A-C). Collectively, these autoradiographic data, when analyzed together with the histological and non-radioactive AMPase activity assays (Fig. 1 and 2), provide evidence for the ability of both tracers, [^18^F]**1** and [^18^F]**2**, to selectively bind to CD73 in human breast cancer samples.

**Fig. 3.**
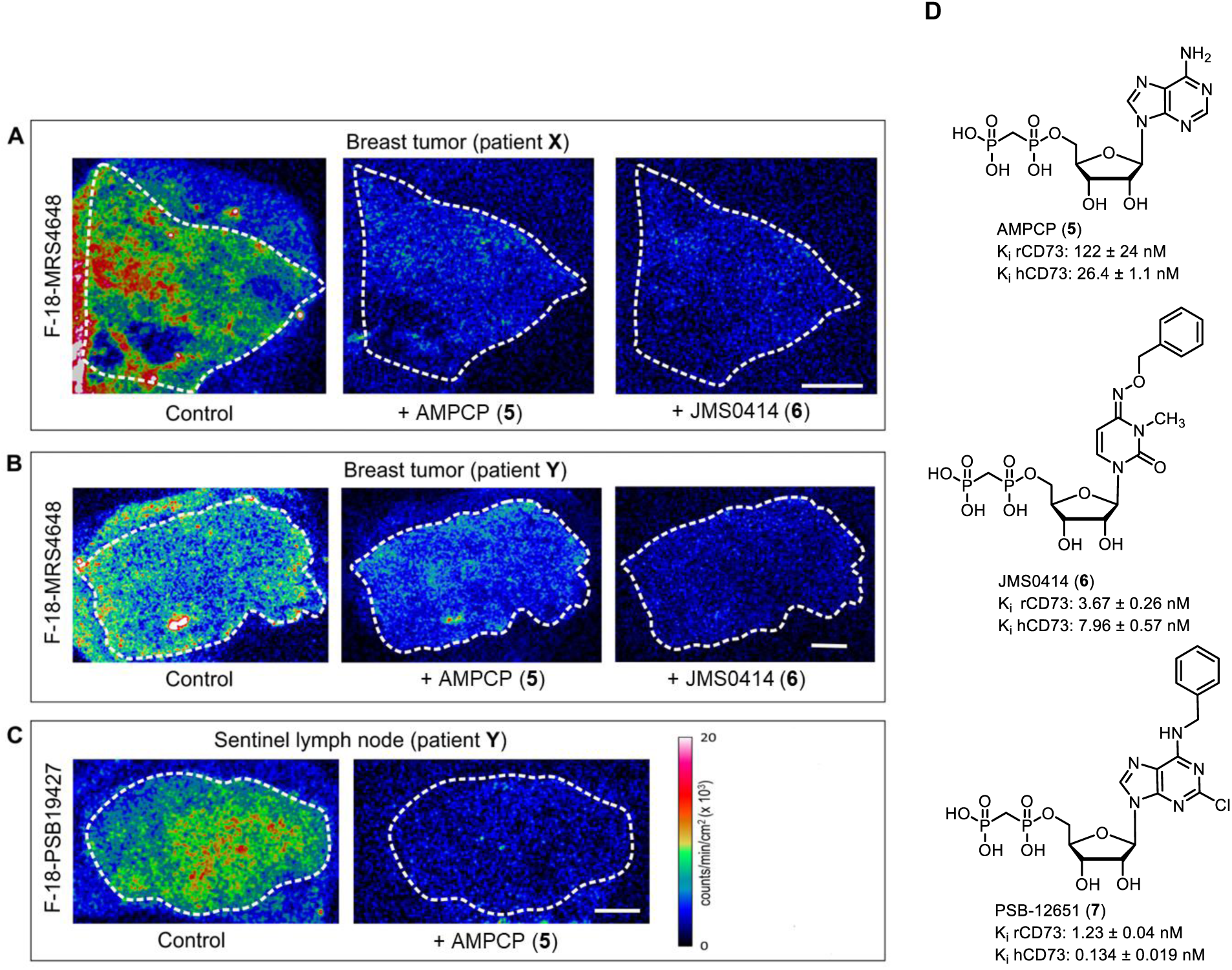
Autoradiographic imaging of ^18^F-labeled CD73 tracer binding to tissue samples of breast tumors and sentinel lymph nodes. Primary breast tumor (A, B) and LN (C) cryosections were incubated with [^18^F]**2** or [^18^F]**1** in the absence (control) and in the presence of unlabeled CD73 inhibitors AMPCP (**5**) or JMS0414 (**6**), as indicated. The samples were processed for autoradiographic analysis, as described in Materials and Methods. The white dotted lines outline the tissue boundaries. (D). Structures of CD73 inhibitors applied as blockers in this study.

### Biodistribution studies of [^18^F]PSB-19427 and [^18^F]MRS-4648 in mice

Biodistribution studies of [^18^F]**1** and [^18^F]**2** were performed in female C57BL/6 WT mice. Dynamic (0-90 min), static (77-90 min), and late (240-260 min) scans were performed in a small animal PET scanner. Each PET scan was combined with computer tomography (CT). In the case of [^18^F]**2**, > 95% of the tracer was excreted within the first 30 minutes via renal (19 %ID) and hepatobiliary (60 %ID, Fig. 4 A-C) routes. After 90 min, most of the tracer was found in the intestine, gallbladder, bladder (urine), and liver, while only 0.3 ± 0.1 %ID/mL of the radiosignal was detected in the blood at that time point (Fig. 4 B, C). Concentrations in the lung, heart, muscle, and brain were also relatively low. Nevertheless, during the first 3 min, the tracer was distributed within the whole body, reaching every tissue (Fig. 4 A).

**Fig. 4:**
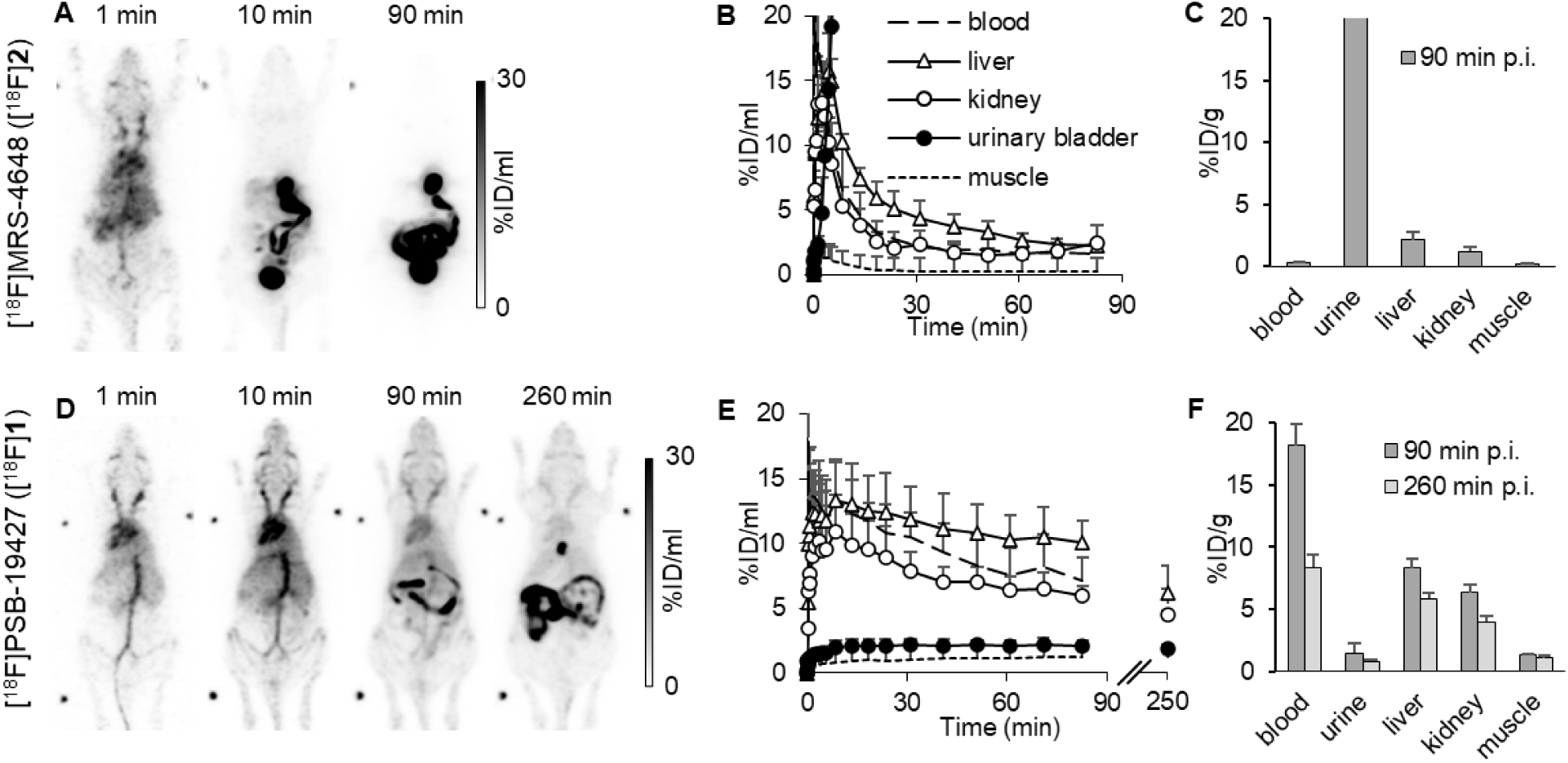
Biodistribution of the investigated CD73 PET tracers. A-C. Biodistribution of [^18^F]MRS-4648 ([^18^F]**2**) in C57BL/6 WT mice. A. PET maximum intensity projections of a representative mouse B. Quantification of tracer distribution (n = 3) C. Sum of ex vivo gamma counter experiments (n = 3). D-F. Biodistribution of [^18^F]PSB-19427 ([^18^F]**1**) in C57BL/6 WT mice. D. Maximum intensity projections of a representative mouse. E. Quantification of tracer distribution (n = 3) F. Sum of ex vivo gamma counter experiments after 90 min (n = 3) and 260 min (n = 3).

The biodistribution of [^18^F]**1** was significantly different from that of [^18^F]**2** (Fig. 4 D-F). The radiotracer [^18^F]**1** was injected into the tail vein, entered the heart, and was subsequently distributed throughout the whole body. The renal excretion route was negligible (2 %ID), with the primary excretion route being hepatobiliary (37 %ID at 90 min p.i.) The tracer showed long retention in blood, while the concentration in muscle was low (Fig. 4 E). Nearly no tracer accumulated in the brain and the urinary bladder, facilitating a favorable signal-to-noise ratio. We prolonged the imaging and performed an additional PET scan after 260 min, to potentially even further increase the signal-to-background ratio. In all investigated residual organs (kidney, lung, heart, liver, and spleen), [^18^F]**1** was present even after 260 min, indicating a favorable biodistribution and high metabolic stability in vivo (Fig. 4 F).

### Evaluation of [^18^F]PSB-19427 ([^18^F]1) and [^18^F]MRS-4648 ([^18^F]2) in a mouse MDA-MB-231 breast cancer model

Given the specificity of [^18^F]**1** and [^18^F]**2** and the promising in vivo biodistribution profile of [^18^F]**1**, we established a small-animal human breast tumor xenograft model in NSG mice subcutaneously implanted with MDA-MB-231 cells. This highly invasive triple-negative breast adenocarcinoma cell line displays a high expression of CD73 and, therefore, offers a suitable model for investigating the role of CD73 expression in TNBC(*36, 37*). For blocking studies, PSB-12651 (**7**, Fig. 5) (*38*) or unlabeled PSB-19427 (**1**) were used.

**Fig. 5:**
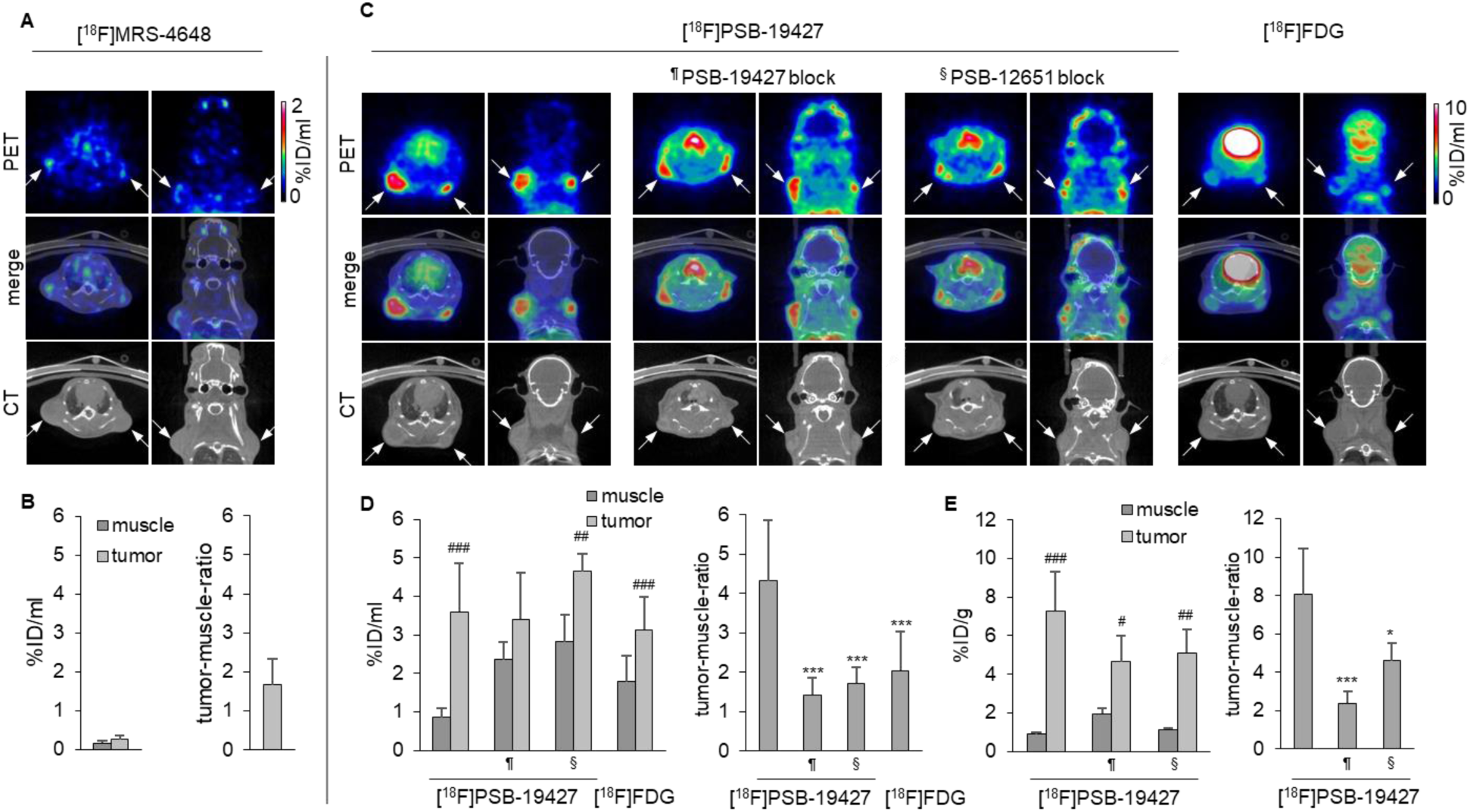
PET imaging data in an MDA-MB-231 tumor mouse model. A. PET images, CT scans, and merged images of [^18^F]**2** in the MDA-MB-231 tumor mouse model 90 min after tracer injection. White arrows mark the tumor positions. B. Quantitative PET analysis of [^18^F]**2** uptake: activity concentration of [^18^F]**2** in muscle and tumor tissues 90 min after tracer injection (left) and respective tumor-to-muscle ratio (right, n = 5 tumors). C. PET-CT images of [^18^F]**1** and [^18^F]FDG in the MDA-MB-231 tumor mouse model, 260 min ([^18^F]**1**), or 75 min ([^18^F]FDG) after injection. From left to right: [^18^F]**1** without blocker, pretreating with unlabeled compound **1** (^¶^PSB-19427) or ^§^PSB-12651 (**7**) 10 min before tracer injection, and [^18^F]FDG. White arrows mark the positions of tumors. D. Quantitative PET analysis of [^18^F]**1** uptake: activity concentration of [^18^F]**1** and [^18^F]FDG in muscle and tumor tissues (left) and tumor-to-muscle ratios (right) after 260 min. [^18^F]PSB-19427: n = 18 tumors, [^18^F]PSB-19427 plus PSB-19427 block: n = 8 tumors, [^18^F]PSB-19427 plus PSB-12651 block: n = 6 tumors, [^18^F]FDG: n = 18 tumors E. Ex vivo analysis of [^18^F]**1** uptake as measured by gamma counter: activity concentrations of [^18^F]**1** in muscle and tumor tissues after 260 min without and with blocking (left) and tumor-to-muscle ratios (right). [^18^F]PSB-19427: n = 6 tumors, [^18^F]PSB-19427 plus PSB-19427 block: n = 6 tumors, [^18^F]PSB-19427 plus PSB-12651 block: n = 5 tumors. # relative to the respective muscle, *relative to [^18^F]**1** without blocker; <0,05; ** <0,01, ***<0,001, unpaired t-test

Similar to the initial radiotracer biodistribution study (Fig. 4 A), [^18^F]MRS-4648 ([^18^F]**2**) distribution was characterized by rapid liver uptake and clearance through both renal and hepatobiliary routes, which was accompanied by very low accumulation in the tumor (Fig. 5 A and B). At the end of the study, 90 min p.i., most of the tracer was detected in the bladder. Also, relevant concentrations were located in the liver, spleen, and kidneys, indicating that the tracer was rapidly excreted.

Unlike [^18^F]**2**, [^18^F]PSB-19427 ([^18^F]**1**) showed elongated blood retention with sufficient accumulation in the tumor tissues. The tumors were visible in the PET scans after 260 min (Fig. 5 C). By pretreating the mice with unlabeled PSB-19427 (**1**) or a structurally different CD73 inhibitor, PSB-12651 (**7**), the tumor-to-muscle ratio was markedly reduced, confirming the high specificity of [^18^F]**1** accumulation in the tumor tissue (Fig. 5 C-E). MDA-MB-231 tumor uptake of [^18^F]PSB-19427 was at 3.60 ± 1.27 %ID/mL, while 0.86 ± 0.24 %ID/mL was found in the muscle, reflecting a low background with a corresponding tumor-to-muscle ratio of 4.32 ± 1.54 after 260 min. Tumor uptake was relatively stable over more than four hours (Fig. S2).

As a next step, we compared the performance of [^18^F]**1** in the MBA-MB-231 xenograft mouse model to [^18^F]FDG PET/CT as a clinical diagnostic approach, which is recommended for the systemic staging (stages IIB-IV) of no special type breast cancer(*39*), albeit its known limited diagnostic accuracy, in particular in lower stages of disease, due to low tracer uptake and detection of false positive findings(*39, 40*). While the MDA-MB-231 tumors are only barely visible by [^18^F]FDG imaging (tumor-to-muscle ratio of 2.04 ± 1.0), the tumors can be clearly identified in the same animals by [^18^F]**1** imaging (tumor-to-muscle ratio of 3.55 ± 1.38 after 90 min, Fig S3. Thus, in the TNBC model using human MDA-MB-231 cells, [^18^F]PSB-19427 ([^18^F]**1**) has a much higher sensitivity and is, therefore, outperforming [^18^F]FDG in PET imaging.

### Imaging of pancreatic cancer by [^18^F]PSB-19427 ([^18^F]1)

As previously mentioned, encouraging clinical results obtained with the nucleotide-derived CD73 inhibitor Quemliclustat have been obtained in the treatment of pancreatic cancer(*41*). Therefore, we expanded our experiments to a human pancreatic cancer (AsPC-1) mouse model. The AsPC-1 cell line shows lower CD73 expression(*42*), is more aggressive and faster-growing and is forming more diffuse tumors compared to the compact and well-defined MDA-MB-231 tumors. In analogy to the results in the MDA-MB-231 tumor model, the pancreatic tumors accumulated the radiotracer [^18^F]PSB-19427, while in the blocked mice, tumors could hardly be delineated on PET images ([^18^F]**1**, Fig. 6 A left). The tumor-to-muscle ratio was 2.49 ± 0.28, while it was reduced to 1.44 ± 0.47, after blocking (Fig. 6B and 6C).

**Fig. 6:**
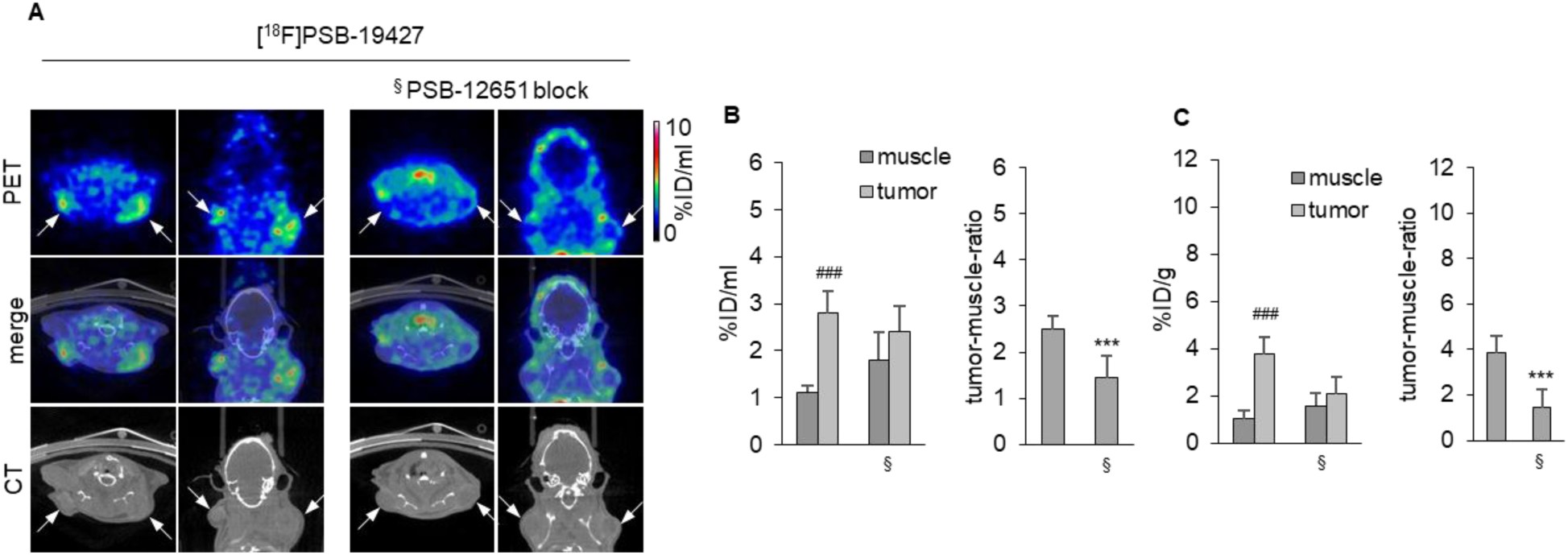
PET/CT imaging and quantitative analysis of [^18^F]PSB-19427 ([^18^F]1) uptake in a pancreatic cancer cell (AsPC-1) xenograft tumor model in mice. A. PET images, CT scans, and merged images of [^18^F]PSB-19427 ([^18^F]**1**) 260 min after tracer injection. Left: no blocker. Right: Pretreatment with PSB-12651 (**7**) 10 min before tracer injection (§). White arrows mark the positions of tumors. B. Quantitative PET analysis of [^18^F]PSB-19427 ([^18^F]**1**) uptake: activity concentration of [^18^F]**1** in muscle and tumor tissues after 260 min without and with blocking (§) (left) and tumor-to-muscle ratios (right). [^18^F]PSB-19427: n = 8 tumors, [^18^F]PSB-19427 plus PSB-12651 block: n = 8 tumors. C. Ex vivo gamma counter analysis of explanted tissues: activity concentration of [^18^F]**1** in muscle and tumor tissues after 260 min without and with (§) blocking (left) and tumor-to-muscle ratios (right). [^18^F]PSB-19427: n = 8 tumors, [^18^F]PSB-19427 plus PSB-12651 block: n = 8 tumors. # relative to the respective muscle, *relative to [^18^F]**1** without blocker; <0,05; ** <0,01, ***<0,001, unpaired t-test

### X-ray co-crystal structure

To further characterize the CD73 interactions of the promising CD73 inhibitor PSB-19427 (**1**), we determined its co-crystal structure (Figure 7). PSB-19427 adopts a binding mode in the closed conformation of CD73 similar to that of other AMPCP derivatives; the large N^6^-substituent interacts with the N-terminal domain. In comparison to the CD73×AMPCP complex in the closed state in crystal form III(*43*) the adenine ring of PSB-19427 is rotated slightly such that the N^6^ atom shifts by about 1.0 Å towards the C-terminal domain (Figure 7B). We also superimposed PSB-12489(*38*), which structurally closely resembles PSB-19427 (**1**) (Figure S4A). The benzyl groups at N^6^ of the two derivatives show a close agreement in their binding modes. It is important to note that the electron density of the N^6^-substituents of **1** is significantly weaker than the rest of the inhibitor structures (Figure S4B). This finding indicates the flexibility of this substituent, which has also been observed previously for most other co-crystal structures of CD73 with N^6^-substituted inhibitors(*19, 20, 38*). Interestingly, the triazole group is better defined in the electron density than the phenyl ring in both chains of the asymmetric unit, and it forms non-polar interactions with Asn186. This interaction may contribute to the compound’s binding properties.

**Fig. 7:**
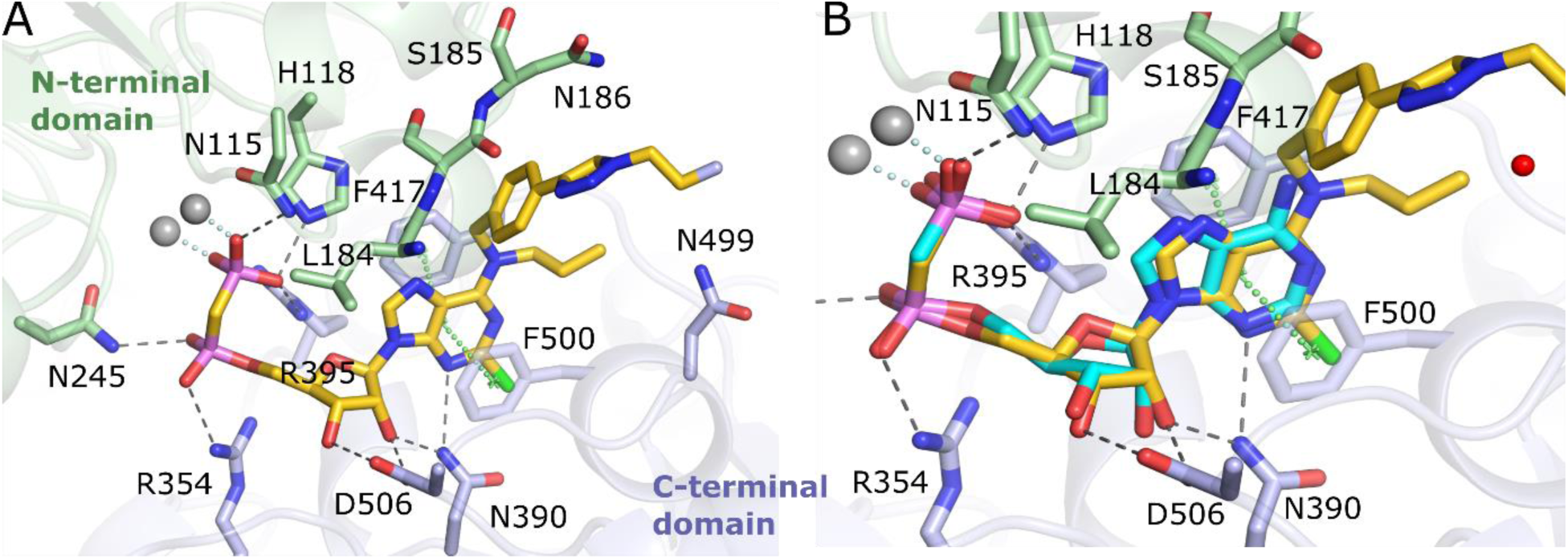
X-ray crystal structure of CD73 in complex with PSB-19427. (**1**). A. Binding mode of PSB-19427 (**1**) in the active site of CD73. Only selected residues interacting with PSB-19427 via polar or hydrophobic contacts are shown. Gray spheres indicate the two zinc ions. B. Superposition of CD73×PSB-19427 (yellow) and CD73×AMPCP (cyan, PDB: 4H2I).

## DISCUSSION

The development of [^18^F] fluorine-labeled CD73 inhibitors as novel PET tracers represents a significant advancement in the imaging of CD73 expression in cancer. While CD73 is ubiquitously expressed throughout the body, its upregulation is strongly correlated with tumor formation (*44*), and the prognostic value of CD73 expression was demonstrated in many different cancer types, including diagnostically challenging TNBC and pancreatic cancer (*14, 15, 45*), but also on immune cells present in the tumor microenvironment or infiltrated into the tumor (*46*). Motivated by the clinical need for a highly specific imaging tracer for such diagnostically challenging types of cancer, we have developed two novel, highly potent, ^18^F-labeled CD73 inhibitors, [^18^F]PSB-19427 ([^18^F]**1,** K*_i_* = 1.02 ± 0.11 nM) and [^18^F]MRS-4648 ([^18^F]**2,** K*_i_* = 0.664 ± 0.089 nM) as potential PET tracers for cancer imaging.

First, PSB-19427 (**1**) and MRS-4648 (**2**) were evaluated in biopsy tissues of breast cancer patients confirming the presence, activity, and tissue-specific distribution of CD73. Subsequently, both derivatives were prepared in their radiolabeled form, starting from the respective precursors **3** or **4** via Huisgen cycloaddition using 18-fluorine labeled 2-fluoroethyl azide providing [^18^F]**1** and [^18^F]**2** in satisfactory radiochemical yield and molar activity, with excellent (>98%) radiochemical purity. Both tracers were stable when formulated for intravenous injection.

Since we observed that tracers [^18^F]**1** and [^18^F]**2** bound selectively to CD73 in human breast cancer samples in autoradiography studies, we proceeded to biodistribution studies in female C57BL/6 WT mice. Despite a certain structural similarity, both radiotracers being AMPCP (**5**) derivatives, the pyrimidine [^18^F]MRS-4648 ([^18^F]**2**) was rapidly excreted, while the purine [^18^F]PSB-19427 ([^18^F]**1**) displayed long retention in blood and was present even after 260 min, indicating a good biodistribution and high metabolic stability in vivo. One possible explanation for the different in vivo behavior could be differences in the compounds’ PPB, 66% for **2** versus >99% for **1**, also reflected by the different radiosignal proportion in the blood (0.3 ± 0.1 %ID/g for [^18^F]**2** versus 18.2 ± 1.7 % ID/g for [^18^F]**1** at 90 min p.i). High PPB is known to protect tracers from metabolic conversion, and increasing this parameter is employed as a common strategy for increasing the circulation time and the bioavailability of radiotracers. While MRS-4648 (**2**) displayed higher in vitro metabolic stability (3% conversion in MLM within 90 min) versus PSB-19427 (**1**, 17% conversion in MLM), it was rapidly excreted in vivo, preventing its accumulation in the MDA-MB-231 xenograft tumor model. In contrast, [^18^F]PSB-19427 ([^18^F]**1**) accumulated sufficiently in the tumor tissues, providing a tumor-to-muscle ratio of 4.3 ± 1.5 after 260 min. Tracer contrast was markedly reduced by applying non-labeled PSB-19427 (**1**) or the structurally different CD73 inhibitor PSB-12651 (**7**). A redistribution of the tracer into the muscle could be observed during the blocking experiments. This might again be explained by the high PPB of the compound, as the high blocker concentration (∼1000 fold higher than the radiotracer) displaced the tracer from the tumor tissue and allowed binding to muscle tissue. Furthermore, a saturation of the CYP enzymes and, thus, different metabolism and excretion, leading to slight differences in biodistribution, is feasible at this blocker concentration. Next, we compared the performance of [^18^F]**1** versus [^18^F]FDG in the MBA-MB-231 xenograft mouse model. While [^18^F]FDG remains a cornerstone of PET imaging, particularly for its ability to detect a wide range of malignancies, its limitations are well-documented. For example, [^18^F]FDG uptake can be nonspecific due to its dependence on glucose metabolism, which is also elevated in inflammatory conditions and non-cancerous tissues. The MDA-MB-231 tumors were only weakly detectable by [^18^F]FDG imaging with a tumor-to-muscle ratio of 2.04. In contrast, in the mice imaged with the tracer [^18^F]**1**, the tumors were clearly visible with a tumor-to-muscle ratio of 4.32 (3.55 after 90 min p.i., Figure S2). Thus, [^18^F]PSB-19427 ([^18^F]**1**) is outperforming [^18^F]FDG in the PET imaging of breast cancer, at least in the employed TNBC mouse model. [^18^F]**1** is specifically targeting CD73, providing a more precise and reliable imaging modality for tumors where CD73 is upregulated. This specificity could reduce false positives and improve the accuracy of tumor detection, particularly in early-stage cancers or in tissues where [^18^F]FDG might yield inconclusive results.

Next, in order to demonstrate its applicability for the imaging of different types of tumors, we employed [^18^F]PSB-19427 ([^18^F]**1**) in PET imaging of a human pancreatic cancer (AsPC-1) mouse model, employing a cancer cell line known for its lower CD73 expression in comparison to MDA-MB-231 cells. Again, [^18^F]PSB-19427 ([^18^F]**1**) displayed a pronounced tumor accumulation with a tumor-to-muscle ratio of 2.49 ± 0.28 after 260 min. The signal was significantly diminished by applying PSB-12651 (**7**) as a CD73 blocker.

CD73 upregulation is strongly correlated to the formation of solid tumors, and therefore, radiolabeled CD73 inhibitors have a high potential for use as pan-PET tracers for tumor imaging. High-resolution methods for solid tumor imaging remain an unmet medical need and, if clinically successful, could save many lives through early cancer detection. The promising results with [^18^F]**1** suggest that CD73-targeted PET imaging could play a crucial role in the diagnosis and staging of cancers with high CD73 expression. Furthermore, the ability of [^18^F]**1** to outperform [^18^F]FDG in specific contexts points to its potential as a preferred imaging agent in certain clinical scenarios, particularly for cancers that are difficult to detect with current methods. Furthermore, due to CD73’s functional relevance and potential as a therapeutic target in immune oncology, the theranostic use of radiolabeled CD73 inhibitors appears promising.

This study is not without limitations. The radiolabeling of [^18^F]PSB-19427 ([^18^F]**1**) through the Huisgen cycloaddition reaction relies on copper catalysis. Developing alternative radiolabeling strategies or modifying the tracer’s chemical structure to enable copper-free labeling could facilitate its transition to clinical settings. Additionally, while the current study focuses on breast and pancreatic cancers, expanding the evaluation of [^18^F]PSB-19427 to other cancer types where CD73 is overexpressed could further validate its utility as a pan-cancer imaging agent.

Moreover, due to the polar structure of the bisphosphonate-bearing nucleotide derivative **1**, the developed PET tracer cannot penetrate the blood-brain barrier and be applied for brain imaging unless administered intrathecally.

In conclusion, the development of [^18^F]PSB-19427 and [^18^F]MRS4648 marks an important step forward in the field of cancer imaging. The high specificity and favorable biodistribution of [^18^F]PSB-19427, in particular, suggest that it could become a valuable tool for the early detection and monitoring of cancers with elevated CD73 expression. Continued research into the optimization and clinical translation of these tracers will be critical in realizing their full potential for the imaging of solid tumors.

## MATERIALS AND METHODS

### Study design

The primary research objective was to design and synthesize an imaging agent to image tumor-specific CD73 expression in living subjects noninvasively via PET. Six to ten animals per group were used to evaluate CD73 expression in xenograft models of breast (MDA-MB-231) and pancreatic (AsPC-1) cancer. All outliers were included in the analysis, and no data were excluded. The authors were not blinded to the results. A minimum of three experimental replicates were recorded for all in vitro data.

Adult, 9 to 13 week old, C57bl/6 (biodistribution study, 19.8±1.3 g) or 7 to 12 weeks old NOD.Cg-*Prkdc^scid^ Il2rg^tm1Wjl^*/SzJ mice (22.3±2.2 g) were obtained from Charles River Laboratories. A total of 43 mice were used for this work

All animal experiments were performed in accordance with the legal requirements of the European Community (Directive 2010/63/EU) and the corresponding German Animal Welfare Law (TierSchG, TierSchVersV) and were approved by the local authorizing agency (State Office for Nature, Environment and Consumer Protection North Rhine-Westphalia).

### Experimental section

#### Chemistry General

All reagents were commercially obtained from various producers (Alfa Aesar, Carbosynth, and Sigma Aldrich) and used without further purification. The purity of all compounds, including starting materials, was more than 95%, as determined using HPLC. Commercial solvents of specific reagent grades were used without additional purification or drying. Analytical thin-layer chromatography was carried out on Sigma-Aldrich® TLC plates and compounds were visualized with UV light at 254 nm. Flash column chromatography (fc): Silica gel 60, 40–64 µm; parentheses include: diameter of the column, length of the column, fraction size, eluent, R_f_ value. Melting point: melting point apparatus Stuart Scientific^®^ SMP 3, uncorrected. IR: IR spectrophotometer FT-ATR-IR (Jasco®). The ^1^H, ^31^P, and ^13^C NMR spectra were recorded using Bruker 400 MHz spectrometer, a DD2 400 MHz or DD2 600 MHz NMR spectrometer (Agilent). DMSO-d_6_, MeOD-d_4_, CDCl_3_ or D_2_O were used as solvents. Shifts are given in ppm relative to the remaining protons of the deuterated solvents used as internal standard (^1^H-, ^13^C-NMR) or using D_3_PO_4_ as external standard (^31^P), the assignments of ^13^C and ^1^H NMR signals were supported by 2D NMR techniques. Purification of final compounds was performed by semi-preparative HPLC (Column: Luna 5 µm C18(2) 100 Å, LC Column 250 x 4.6 mm). Eluent: 10 mM triethylammonium acetate buffer - CH_3_CN from 80:20 to 20:80 in 40 min, with a flow rate of 5 mL/min. Purities of all tested compounds were ≥95%, as estimated by analytical HPLC: Equipment: UV-detector: UltiMate 3000 variable Wavelength Detector; autosampler: UltiMate 3000; pump: Ultimate 3000; degasser: Ultimate 3000: data acquisition: Chromeleon Client 8.0.0 (Dionex Corpor.). Method: flow rate: 1.00 mL/min; injection volume: 5.0 µL; <u>Method A</u>: Eluent: 5 mM triethylammonium phosphate monobasic solution - CH_3_CN from 100:0 to 50:50 in 20 min, then triethylammonium phosphate monobasic solution - CH_3_CN to 100:0 in 5 min with a flow rate of 1 mL/min (Column: Zorbax SB-Aq 5 µm analytical column, 50 X 4.6 mm; Agilent Technologies, Inc). <u>Method B</u>: Eluent: 5 mM triethylammonium phosphate monobasic solution - CH_3_CN from 90:10 to 0:100 in 20 min, then triethylammonium phosphate monobasic solution - CH_3_CN from 0:100 to 90:10 in 5 min with a flow rate of 1 mL/min (Column: Zorbax SB-Aq 5 µm analytical column, 150 X 4.6 mm; Agilent Technologies, Inc). <u>Method C</u>: Eluent: 5 mM triethylammonium phosphate monobasic solution - CH_3_CN from 80:20 to 20:80 in 20 min, then triethylammonium phosphate monobasic solution - CH_3_CN from 20:80 to 80:20 in 10 min with a flow rate of 1 mL/min (Column: Zorbax SB-Aq 5 µm analytical column, 150 X 4.6 mm; Agilent Technologies, Inc). Peaks were detected by UV absorption (210 nm) using a UV/VIS detector. Low-resolution mass spectrometry was performed with a JEOL SX102 spectrometer with 6-kV Xe atoms following desorption from a glycerol matrix or on an Agilent LC/MS 1100 MSD, with a Waters (Milford, MA) Atlantis C18 column. High-resolution mass spectroscopic (HRMS) measurements were performed on a proteomics optimized Q-TOF-2 (Micromass-Waters) using external calibration with polyalanine or a MicroTof (Bruker Daltronics, Bremen), calibration with sodium formate clusters before measurement. For lyophilization, a freeze dryer (Labconco FreeZone 4.5) was used.

#### [((2*R*,3*S*,4*R*,5*R*)-5-{2-Chloro-6-[(4-ethynylbenzyl)(propyl)amino]-9*H*-purin-9-yl}(*11*)-3,4-dihydroxytetrahydrofuran-2-yl)methoxy] (phosphonomethyl)-phosphonic acid (3)

(2*R*,3*R*,4*S*,5*R*)-2-(2-Chloro-6-((4-ethynylbenzyl)(propyl)amino)-9*H*-purin-9-yl)-5- (hydroxymethyl)tetrahydrofuran-3,4-diol (**11**, 0.1 g, 0.22 mmol, 1.0 eq.) was dissolved in trimethyl phosphate (2 mL) and stirred at 0-4 °C. A solution of methylenebis(phosphonic dichloride) (0.27 g, 1.09 mmol. 5.0 eq.) in trimethyl phosphate (3 mL), cooled to 0-4 °C was added. The reaction mixture was stirred at 0-4 °C and samples were withdrawn at 15 min intervals for TLC to control for the disappearance of nucleosides. After 30 min, when the nucleoside had completely reacted, 20 mL of cold 0.5 M aqueous TEAC solution (pH 7.4-7.6) was added and the solution was stirred at 0 °C for 15 min followed by stirring at room temperature for 1 h. Trimethyl phosphate was extracted using 2 x 250 mL of *tert*-butyl methyl ether, and the aqueous layer was lyophilized. The crude product was then purified by RP-HPLC (0-50% H_3_CCN/50 mM NH_4_HCO_3_ buffer within 20 min, 20 mL/min) and the appropriate fractions were pooled. The product was obtained as a white solid after lyophilization (0.11 g, 89%). Purity determined by HPLC-UV (254 nm)-ESI-MS: 84%. mp: 141 °C. ^1^H NMR (600 MHz, D_2_O) *δ* 8.37 (s, 1H, 8-C*H*), 7.30 (br s, 2H, 3-CH_phenyl_, 5-CH_phenyl_), 7.18 (d, 2H, *J* = 7.78 Hz, 2-CH_phenyl_, 6-CH_phenyl_), 6.01 (d, 1H, *J* = 5.36 Hz, 1ʹ-C*H*), 5.31 (br s, 2H, NC*H_2_*), 4.68 (t, 1H, *J* = 5.20 Hz, 2ʹ-C*H*), 4.51 (d, 1H, *J* = 4.70 Hz, 3ʹ-C*H*), 4.36 (d, 1H, *J* = 3.79 Hz, 4ʹ-C*H*), 4.16 (t, 2H, *J* = 4.24 Hz, 5ʹ-C*H*_2_), 3.92 (br s, 2H, NC*H_2_*), 3.43 (s, 1H, CC*H*), 2.19 (t, 2H, *J* = 19.68 Hz, PC*H_2_*P), 1.62 (br s, 2H, C*H_2_*CH_3_), 0.84 (t, 3H, *J* = 7.55 Hz, C*H_3_*). ^13^C NMR (126 MHz, D_2_O) *δ* 167.5, 157.6, 156.4, 154.1, 141.3, 141.1, 134.9, 130.3, 130.1, 125.7, 123.1, 120.8, 89.8, 86.6, 85.9, 81.0, 77.2, 73.0, 66.4, 53.5, 30.4, 23.86, 13.1. ^31^P NMR (202 MHz, D_2_O) *δ* 18.95 (d, 1P, *J* = 9.50 Hz, β-*P*) 14.95 (d, 1P, *J* = 9.28 Hz, α-*P*). Exact mass (ESI): m/z = calcd. for C_23_H_29_ClN_5_O_9_P_2_ 616.1129, found 616.1130 [M+H]^+^ and calcd. for C_23_H_27_ClN_5_O_9_P_2_ 614.0973, found 614.0991 [M-H]^-^.

#### {[{[(2*R*,3*S*,4*R*,5*R*)-5-(2-Chloro-6-{4-[1-(2-fluoroethyl)-1*H*-1,2,3-triazol-4-yl) benzyl](propyl)amino}-9*H*-purin-9-yl)-3,4-dihydroxytetrahydrofuran-2-yl]methoxy{(hydroxy)phosphoryl]methyl}phosphonic acid (1, PSB-19427)

Compound **3** (15.0 mg, 0.02 mmol, 1.0 eq) was dissolved in THF/H_2_O/*t*-BuOH (3:1:1, 0.5 mL). Crude 2-fluoroethylazide 10 (50.0 μl) was added. Additionally, 1 M sodium ascorbate in H_2_O (0.03 mL, 0.03 mmol, 1.2 eq) was added. Finally, a premixed solution of CuSO_4_ (1.0 mg, 0.009 mmol, 0.3 eq) and TBTA (4.0 mg, 0.003 mmol, 0.3 eq) in THF/H_2_O/*t*-BuOH (3:1:1, 0.5 mL) was added to the reaction mixture. The reaction was stirred in the dark under argon at rt. After one night of stirring the reaction mixture was evaporated and directly purified by RP-HPLC (0-70% MeCN/50mM NH_4_HCO_3_ buffer within 20 min, 20 mL/min). Appropriate fractions were pooled and lyophilized overnight to obtain the final product as a white solid (0.003 g, 17.5%). Purity determined by HPLC-UV (254 nm)-ESI-MS: 99%. mp. 169 °C. ^1^H NMR (600 MHz, D_2_O) *δ* 8.40 (s, 1H, C*H*_triazole_), 8.27 (s, 1H, 8-C*H*_purine_), 7.67 (d, 2H, *J* = 6.91 Hz, 3-C*H*_phenyl_, 5-C*H*_phenyl_,), 7.36 (d, 2H, *J* = 7.90 Hz, 2-CH_phenyl_, 6-CH_phenyl_), 6.02 (d, 1H, *J* = 4.74 Hz, 1ʹ-C*H*), 4.93-4.85 (m, 2H, NC*H_2_*), 4.77 (m, 2H, overlapping with H_2_O: NC*H*_2_), 4.73 (s, 1H, 2ʹ-C*H*), 4.53 (s, 1H, 3ʹ-C*H*), 4.38 (s, 1H, 4ʹ-C*H*), 4.18 (br s, 2H, 5ʹ-C*H*_2_), 3.15 (m, 2H, NC*H*_2_), 2.18 (br s, 2H, PCH_2_P), 1.67 (m, 2H, C*H*_2_F), 1.34 (m, 2H, C*H_2_*CH_3_), 0.91 (m, 3H, C*H_3_*). ^13^C NMR (126 MHz, D_2_O) *δ* 159.1, 156.5, 154.1, 152.6, 143.9, 141.0, 131.3, 131.0, 128.6, 125.4, 120.8, 89.7, 86.7, 85.5, 77.1, 73.1, 66.5, 55.5, 53.8, 53.6, 49.5, 28.1, 15.6. ^31^P NMR (243 MHz, D_2_ O) δ 19.43, 14.48. ^19^F NMR (471 MHz, D_2_O) δ -223.05.Exact mass (ESI): m/z = calcd. for C_25_H_33_ClFN_8_O_9_P_2_ 705.1518, found 705.1486 [M+H]^+^ and calcd. for C_25_H_31_ClFN_8_O_9_P_2_ 703.1362, found 703.1335 [M-H]^-^.

#### {[{[(2*R*,3*S*,4*R*,5*R*)-5-(-4-{[(4-Ethynylbenzyl)oxy]imino}-3-methyl-2-oxo-3,4-dihydropyrimidin-1(2*H*)-yl)-3,4-dihydroxytetrahydrofuran-2-yl]methoxy}(hydroxy)phosphoryl]methyl}phosphonic acid (4)

1-[(2*R*,3*R*,4*S*,5*R*)-3,4-Dihydroxy-5-(hydroxymethyl)tetrahydrofuran-2-yl]-4-{[(4-ethynylbenzyl)oxy]imino}-3-methyl-3,4-dihydropyrimidin-2(1*H*)-one (**15**, 208 mg, 0.38 mmol, 1.0 eq.) was suspended in trimethyl phosphate (2 mL, 0 °C), and methylenebis(phosphonic dichloride) (143 mg, 0.57 mmol, 1.5 eq.) in trimethyl phosphate (2 mL, 0 °C) was added. The mixture was stirred at 0 °C (controlled by LC-MS) for 85 min, and triethylammonium acetate buffer (TEAA, 0.1 M in water, pH = 7, 40 mL) was added. The solution was neutralized by the addition of triethylamine and was then lyophilized. The compound was then pre-purified by RP-HPLC-A and purified by RP-HPLC-B. Suitable fractions were pooled and lyophilized to obtain the final product as a glassy colorless solid (1.17 eq. Et_3_N-salt [663.44 g/mol], 46 mg, 69 µmol, 13 %). *R_t_* (RP-HPLC-A): 27.7-29.5 min: solvent A: CH_3_CN, solvent B: 0.1 M TEAA, 0-2 min: 25 % solvent A, 3 mL/min → 5 mL/min, 2-30 min: 40 % solvent A, 30-40 min: 40 % solvent A, 40-41 min: 40 % → 100 % solvent A, 41-51 min: 100 % solvent A, 51-52 min: 100 % to 25 % solvent A, 52-59 min: 25 % solvent A, 59-60 min: 25 % solvent A, 5 mL/min to 0.1 mL/min, Injection volume: 300 µL. *R_t_* (RP-HPLC-B): 31.0-33.0 min: solvent A: CH_3_CN, solvent B: 0.1 M TEAA, 0-2 min: 25 % solvent A, 3 mL/min to 5 mL/min, 2-50 min: 40 % solvent A, 50-52 min: 40 % to 25 % solvent A, 52-59 min: 25 % solvent A, 59-60 min: 25 % solvent A, 5 mL/min to 0.1 mL/min, Injection volume : 400-600 µL. Purity determined by HPLC (210 nm) >99 % (*R_t_* = 12.2 min). Exact mass (ESI): m/z = calcd. for C_20_H_24_N_3_O_11_P_2_ 544.0892, found 544.0862 [M-H]^-^.^1^H NMR (D_2_O, 400 MHz): δ (ppm) = 7.48 (d, *J* = 8.2 Hz, 2H, 3-C*H*_phenyl_ and 5-C*H*_phenyl_), 7.37 (d, *J* = 8.2 Hz, 2H, 2-C*H*_phenyl_ and 6-C*H*_phenyl_), 7.32 (d, *J* = 8.3 Hz, 1H, 6-C*H*_pyrimidine_), 6.38 (d, *J* = 8.3 Hz, 1H, 5-C*H*_pyrimidine_), 5.89 (d, *J* = 4.8 Hz, 1H, 1ꞌ-C*H*), 4.97 (s, 2H, C*H*_2-benzyl_), 4.34-4.25 (m, 2H, 2ꞌ-C*H* and 3ꞌ-C*H*), 4.22-4.19 (m, 1H, 4ꞌ-C*H*), 4.18-4.09 (m, 2H, 5ꞌ-C*H*_2_), 3.52 (s, 1H, C*H*_ethynyl_), 3.18 (q, *J* = 7.3 Hz, 7H, C*H*_2 NEt3_), 3.11 (s, *3*H, C*H*_3 pyrimidine_), 2.32 (t, *J* = 19.4 Hz, 2H, PC*H*_2_P), 1.26 (t, *J* = 7.3 Hz, 10.5H, C*H*_2 NEt3_). ^13^C NMR (D_2_O, 101 MHz): δ (ppm) = 153.6 (*C*=N_pyrimidine_), 151.0 (*C*=O_pyrimidine_), 138.0 (*C*-1_phenyl_), 133.3 (*C*-6_pyrimidine_), 132.26 (2C, *C*-3_phenyl_ and *C*-5_phenyl_), 128.8 (2C, *C*-2_phenyl_ and *C*-6_phenyl_), 121.3 (*C*-4_phenyl_), 93.8 (*C*-5_pyrimidine_), 88.8 (*C*-1ꞌ), 83.7 (*C*q_ethynyl_), 83.0 (d, *J* = 7.0 Hz, *C*-4ꞌ), 78.7 (*C*H_ethynyl_), 75.2 (*C*H_2-benzyl_), 73.2 (*C*-2ꞌ), 69.8 (*C*-3ꞌ), 63.5 (bs, *C*-5ꞌ), 46.7 (3.5C, *C*H_2 NEt3_), 29.3 (CH_3_), 26.7 (t, *J* = 127.5 Hz, P*C*H_2_P), 8.3 (3.5C, *C*H_3 NEt3_). ^31^P NMR (D_2_O/D_3_PO_4_, 162 MHz): δ (ppm) = 17.8 (bs, 2P).

#### [((2*R*,3*S*,4*R*,5*R*)-5-{4-[({4-[1-(2-Fluoroethyl)-1*H*-1,2,3-triazol-4-yl]benzyl}oxy)-imino]-3-methyl-2-oxo-3,4-dihydropyrimidin-1(2*H*)-yl}-3,4-dihydroxytetrahydro-furan-2-yl)methoxy]-(phosphonomethyl)phosphonic acid (MRS-4648 (2))

Compound **4** (51 mg, 0.13 mmol, 1.0 eq.) was dissolved in trimethyl phosphate (3 mL), and the solution was cooled to 0 °C. Methylenebis(phosphonic dichloride) (50 mg, 0.20 mmol, 1.5 eq.) was added, and the mixture was stirred at 0 °C for 50 min (controlled by LC-MS). Triethylammonium acetate (1 M in water, pH = 7, 1 mL) was added, and the solution was stirred at 0 °C for 30 min and for 1 h at room temperature. The crude product was then purified *via* semi-preparative RP-HPLC (RP-HPLC-J). Fractions containing the product were pooled and lyophilized to obtain the final product as a colorless solid (41 mg, 0.041 mmol, 32 % [M = 998.08 g/mol, 1.0 eq. acetic acid and 3.0 eq. Et_3_N-salt]). *R_t_* (RP-HPLC-J): 37.5-39 min: solvent A: CH_3_CN, solvent B: 0.1 M TEAA, 0-2 min: 15 % solvent A, 3 mL/min → 5 mL/min, 2-30 min: 15 % → 30 % solvent A, 30-35 min: 30 % → 100 % solvent A, 35-40 min: 100 % solvent A, 40-45 min: 100 % → 15 % solvent A, 45-55 min: 15 % solvent A, 55-57 min: 15 % solvent A, 5 mL/min → 0.1 mL/min, Injection volume: 800 µL. column: Agilent technologies, Polaris^TM^ C18-A, 250x21.2 mm, 5 µm, precolumn: Agilent technologies, Pursuit^TM^ C18, 30x21.2 mm, 5 µm. Purity (HPLC-A): 99 % (*R_t_* = 11.3 min). HRMS (ESI): m/z = calcd. for C_22_H_28_FN_6_O_11_P_2_ 633.1273, found 633.1261 [M-H]^-^. ^1^H NMR (D_2_O, 400 MHz): δ (ppm) = 8.38-8.35 (m, 1H, C*H*_triazole_), 7.79 (dd, *J* = 8.2, 2.2 Hz, 2H, 3-C*H*_phenyl_, 5-C*H*_phenyl_), 7.58-7.50 (m, 2H, 2-C*H*_phenyl_, 6-C*H*_phenyl_), 7.32 (dd, *J* = 8.5, 1.1 Hz, 1H, 6-C*H*_pyrimidine_), 6.44 (dd, *J* = 8.4, 1.5 Hz, 1H, 5-C*H*_pyrimidine_), 5.92 (d, *J* = 5.0 Hz, 1H, 1ꞌ-C*H*), 5.03 (s, 2H, C*H*_2-benzyl_), 4.95 (d, *J* = 4.6 Hz, 1H, CH_2_C*H*HF), 4.87-4.81 (m, 2H, CH_2_CH*H*F, C*H*HCH_2_F), 4.77 (m, 1H, CH*H*CH_2_F [below water signal]), 4.35-4.28 (m, 2H, 2ꞌ-C*H*, 3ꞌ-C*H*), 4.22-4.17 (m, 1H, 4ꞌ-C*H*), 4.10 (dd, *J* = 5.2, 2.9 Hz, 2H, 4ꞌ-C*H*_2_), 3.18 (q, *J* = 7.5 Hz, 21H, C*H*_3 pyrimidine_, C*H*_3 NEt3_), 2.17 (td, *J* = 19.8, 1.4 Hz, 2H, PC*H*_2_P), 1.92 (s, 3H, C*H*_3 acetyl_), 1.26 (t, *J* = 7.4 Hz, 27H, C*H*_2 NEt3_). ^13^C NMR (D_2_O, 101 MHz): δ (ppm) = 181.2 (1C, *C*=O_acetyl_), 153.8 (1C, *C*=O_pyrimidine_), 151.2 (1C, *C*=N_pyrimidine_), 147.4 (1C, *C*-1_triazole_), 137.6 (1C, *C*-1_phenyl_), 132.7 (1C, *C*-6_pyrimidine_), 129.6 (2C, *C*-3_phenyl_, *C*-5_phenyl_), 129.4 (1C, *C*-4_phenyl_), 125.8 (2C, *C*-2_phenyl_, *C*-6_phenyl_), 122.9 (1C, *C*-2_triazole_), 94.1 (1C, *C*-5_pyrimidine_), 88.5 (1C, *C*-1ꞌ), 83.2 (d, *J* = 7.9 Hz, 1C, *C*-4ꞌ), 82.2 (d, *J* = 167.0 Hz, 1C, *C*H_2_F), 75.2 (1C, *C*H_2-benzyl_), 73.1 (1C, *C*-2ꞌ), 70.0 (1C, *C*-3ꞌ), 63.7 (1C, *C*-5ꞌ), 50.9 (d, *J* = 19.4 Hz, 1C, *C*H_2_CH_2_F), 46.8 (9C, 1C, *C*H_2 NEt3_), 29.3 (1C, *C*H_3 pyrimidine_), 27.5 (m, 1C, P*C*H_2_P), 23.2 (1C, *C*H_3 acetyl_), 8.3 (9C, *C*H_3 NEt3_). ^19^F NMR (D_2_O, 376 MHz): δ (ppm) = -222.3 (tt, *J* = 46.4, 28.2 Hz, 1F). ^31^P NMR (D_2_O/D_3_PO_4_): δ (ppm) = 18.3 (d, *J* = 9.7 Hz, 1P, α-*P*), 14.7 (d, *J* = 9.8 Hz, 1P, β-*P*).

### Radiochemistry

#### General Methods

The first step of the radiosynthesis was carried out on a modified PET tracer radio synthesizer (TRACERLab Fx_FDG_, GE Healthcare). The recorded data was processed by the TRACERLab Fx software (GE Healthcare). Separation and purification of the radiolabeled compounds were performed on the semipreparative radio-HPLC system A: K-500 and K-501 pump, K-2000 UV detector (Herbert Knauer GmbH), NaI(TI) Scintibloc 51 SP51 γ-detector (Crismatec) and an ACE 5 AQ column (250 mm × 10 mm). Method A started with a linear gradient from 10% to 90% CH_3_CN in water (0.1% TFA) over 30 min, holding for 5 min and followed by a linear gradient from 90% to 10% CH_3_CN in water (0.1% TFA) over 5 min, with l = 254 nm and a flow rate of 5.0 mL min^-1^. Radiochemical purities and molar activities were determined using the analytical radio-HPLC system B: Two Smartline 1000 pumps and a Smartline UV detector 2500 (Herbert Knauer GmbH), a GabiStar γ-detector (Raytest Isotopenmessgeräte GmbH) and a Nucleosil 100-5 C-18 column (250 mm × 4 mm). Method B started with a linear gradient from 10% to 100% CH_3_CN in water (0.1% TFA) over 15 min, holding for 3 min followed by a linear gradient from 100% to 10% CH_3_CN in water (0.1% TFA) over 2 min, with l = 254 nm and a flow rate of 1.0 mL min^-1^. The recorded data of both HPLC-systems were processed by the GINA Star software (Raytest Isotopenmessgeräte GmbH). No-carrier-added aqueous [^18^F]fluoride was produced on a RDS 111e cyclotron (CTI-Siemens) by irradiation of a water target (2.8 mL) using 10 MeV proton beams on 97.0% enriched [^18^O]H_2_O by the ^18^O(p,n)^18^F nuclear reaction.

#### [^18^F]-[((2*R*,3*S*,4*R*,5*R*)-5-{2-chloro-6-[(4-{1-[2-(fluoro)ethyl]-1*H*-1,2,3-triazol-4-yl}benzyl)(propyl)amino]-9*H*-purin-9-yl}-3,4-dihydroxytetrahydrofuran-2-yl)methoxy]methylenebisphosphonic acid ([^18^F]PSB-19427, [^18^F]1)

In a computer controlled TRACERLab Fx_FDG_ Synthesizer a batch of aqueous [^18^F]fluoride (3.1 – 5.2 GBq) from the cyclotron target was passed through an anion exchange resin (pre-conditioned Sep-Pak^®^ Light QMA cartridge with carbonate counter-ion). [^18^F]fluoride was eluted from the resin with a mixture of 1 M K_2_CO_3_ (aqueous, 40 µL), water for injection (WFI, 200 µL), and acetonitrile (800 µL, DNA-grade) containing Kryptofix^®^2.2.2 (K_2.2.2_, 20 mg, 53 µmol) in the reactor. Subsequently, the aqueous K(K_2.2.2_)[^18^F]fluoride solution was carefully evaporated to dryness *in vacuo*. An amount of precursor compound 2-azidoethyl 4-methylbenzenesulfonate (20 mg, 83 µmol) in acetonitrile (DNA-grade, 500 µL) was added and the mixture was heated at 110°C for 3 min. Meanwhile, the labeled 1-azido-2-[^18^F]fluoroethane was distilled from the reactor into an ice-cooled 10 mL flask that contained a mixture of **3** (5.0 mg, 8.1 µmol) in DMF (300 µL), CuSO_4_·5H_2_O (40 mg, 160 µmol) in HEPES buffer (pH: 5.7, 100 µL) and sodium ascorbate (63 mg, 318 µmol) in HEPES buffer (pH: 5.7, 100 µL). After 30 min stirring at 60°C, the mixture was passed through a PTFE sterile filter (0.2 µm). The filter was rinsed with DMF (500 µL) and then with WFI (500 µL). The combined filtrate and rinsing solutions were purified by gradient-radio-HPLC system A (method A). The product fraction of compound [^18^F]PSB-19427 (retention time t*_R_*([^18^F]PSB-19427)= 12.2 min) was collected in a flask pretreated with Sigmacote^®^ and solution was evaporated to dryness *in vacuo*. The residue was redissolved in WFI/EtOH (1 mL, 9:1 v/v). Product compound [^18^F]PSB-19427 was obtained in an overall radiochemical yield of 21.7 ± 3.5% (decay-corrected, based on cyclotron-derived [^18^F]F-ions, n = 21) in 119 ± 10 min from the end of radionuclide production. [^18^F]PSB-19427 was isolated in radiochemical purities of >99% with molar activities in the range of 2.3 – 54.3 GBq/µmol at the end of the synthesis. Radiochemical purities and molar activities of [^18^F]PSB-19427 (retention time t*_R_*([^18^F]PSB-19427)= 9.4 min) were determined by analytical radio-HPLC B (method B). *R_t_* (Radio HPLC-A): 12.2 min. Rcy: 21.7 ± 3.5 %. Rcp (Radio HPLC-B): >99 % (*R_t_* = 9.4 min).

Molar activity: 2.3 – 54.3 GBq/µmol. Log *D*_7.4_: -0.12 ± 0.03. PPB: >99 %. Mouse serum: stable over 90 min. Human serum: stable over 90 min.

#### [^18^F]-{[(2*R*,3*S*,4*R*,5*R*)-5-(4-{[(4-{1-[2-(fluoro)ethyl]-1*H*-1,2,3-triazol-4-yl}benzyl)oxy]imino}-3-methyl-2-oxo-3,4-dihydropyrimidin-1(2*H*)-yl)-3,4-dihydroxytetrahydrofuran-2-yl]methoxy}methylenebisphosphonic acid ([^18^F]MRS-4648, [^18^F]2)

In a computer-controlled TRACERLab Fx_FDG_ Synthesizer a batch of aqueous [^18^F]fluoride (3.8 – 5.4 GBq) from the cyclotron target was passed through an anion exchange resin (pre-conditioned Sep-Pak^®^ Light QMA cartridge with carbonate counter-ion). [^18^F]fluoride was eluted from the resin with a mixture of 1 M K_2_CO_3_ (aqueous, 40 µL), water for injection (WFI, 200 µL), and acetonitrile (800 µL, DNA-grade) containing Kryptofix^®^2.2.2 (K_2.2.2_, 20 mg, 53 µmol) in the reactor. Subsequently, the aqueous K(K_2.2.2_)[^18^F]fluoride solution was carefully evaporated to dryness *in vacuo*. An amount of precursor compound 2-azidoethyl 4-methylbenzenesulfonate (20 mg, 83 µmol) in acetonitrile (DNA-grade, 500 µL) was added and the mixture was heated at 110°C for 3 min. Meanwhile, the labeled 1-azido-2-[^18^F]fluoroethane was distilled from the reactor into an ice-cooled 10 mL flask that contained a mixture of **4** (5.0 mg, 8.1 µmol) in DMF (300 µL), CuSO_4_·5H_2_O (40 mg, 160 µmol) in HEPES buffer (pH: 5.7, 100 µL) and sodium ascorbate (63 mg, 318 µmol) in HEPES buffer (pH: 5.7, 100 µL). After 30 min stirring at 60°C, the mixture was passed through a PTFE sterile filter (0.2 µm). The filter was rinsed with DMF (500 µL) and then with WFI (500 µL). The combined filtrate and rinsing solutions were purified by gradient-radio-HPLC system A (method A). The product fraction of compound [^18^F]MRS-4648 (retention time t*_R_*([^18^F]MRS-4648)= 8.0 min) was collected in a flask pretreated with Sigmacote^®^ and solution was evaporated to dryness *in vacuo*. The residue was redissolved in WFI/EtOH (1 mL, 9:1 v/v). Product compound [^18^F]MRS-4648 was obtained in an overall radiochemical yield of 12.9 ± 3.0% (decay-corrected, based on cyclotron-derived [^18^F]F-ions, n = 8) in 115 ± 15 min from the end of radionuclide production. [^18^F]MRS-4648 was isolated in radiochemical purities of 98 ±1.9% with molar activities in the range of 0.4 – 6.3 GBq/µmol at the end of the synthesis. Radiochemical purities and molar activities of [^18^F]MRS-4648 (retention time t*_R_*([^18^F]MRS-4648)= 8.5 min) were determined by analytical radio-HPLC B (method B). *R_t_* (Radio HPLC-A): 7.9 min. Rcy: 12.9 ± 3.0 %. Rcp (Radio HPLC-B): 98 ±1.9 % (*R_t_* = 8.5 min). Molar activity: 0.4 – 6.3 GBq/µmol. Log *D*_7.4_: 0.74 ± 0.29. PPB: 66.5 %. Mouse serum: stable over 90 min. Human serum: stable over 90 min.

#### Crystal structure analysis

For crystallization of CD73×PSB-19427 our construct 8.01His of human CD73 was used(*26*). This construct is expressed from a derivative of the pHLsec vector, in which the signal peptide of this vector was replaced with the native CD73 signal peptide. The protein was expressed in HEK293S cells in adherent culture in roller bottles. The protein was purified by two chromatography steps. First, the supernatant of the cells was concentrated to 100 mL by ultrafiltration. During ultrafiltration, the culture medium was replaced by a buffer consisting of 50 mM Tris pH 8.0 and 400 mM NaCl. Before application to a 1 mL HisTrap HP column, 4 mL elution buffer (50 mM Tris pH 8.0, 400 mM NaCl, 500 mM imidazole) was added to the 100 mL ultrafiltration concentrate. The HisTrap column was equilibrated with the washing buffer (50 mM Tris pH 8.0, 400 mM NaCl, 20 mM imidazole) before applying the protein solution. After washing the bound protein with 40 mL washing buffer, the protein was eluted with a gradient of 50 mL from washing buffer to the elution buffer. 10 μl 100 mM was added to each fraction tube (fraction sample volume 1 mL) before starting the elution. The pooled fractions were applied to a Superdex 200 16/60 gel filtration column and eluted with the gel filtration buffer consisting of 50 M Tris pH 8.0 and 100 mM NaCl. For crystallization, vapor diffusion trials were set up at 19 °C. 1 µL of the protein solution containing 1 mM of PSB-19427 and 100 µM of ZnCl_2_ were mixed with an equal amount of reservoir buffer solution to give concentrations listed in Table S1. Crystals were stepwise transferred to a cryo buffer composed as reported in Table S1 and finally flash frozen in liquid nitrogen.

X-ray data collection was carried out at 100 K on beamline 14.1 of the Berlin Synchrotron (BESSY, Berlin, Germany) equipped with a PILATUS 6M detector (Dectris, Baden, Switzerland). The diffraction data were integrated with XDS(*47*) via the XDSAPP(*48*) GUI and scaled with StarAniso due to the strong anisotropic diffraction of the crystals. Coordinates of human CD73 in crystal form IV (pdb code: 6ye1)(*25*) were taken as the starting model for automatic refinement with buster and pipedream. COOT(*49*) was used for model building. The two domains of residues 26-333 and 334-551 were used as TLS groups. Structures were validated by MOLPROBITY(*50*). Coordinate files and restraint dictionaries for inhibitors were generated using the GRADE web server (http://grade.globalphasing.org/). Polder omit maps(*51*) were calculated with PHENIX(*52*). Relevant crystallographic parameters of the structures are listed in Table S1. Figures were prepared using PYMOL (http://pymol.org).

#### Stability in mouse liver microsomes (MLM)

An aqueous buffer of 75 mM PBS, 12.5 mM MgCl_2_, 0.6 mM NADPH, 1 mg/mL mouse liver microsomes and 5 µM of the respective compound was prepared. 200 µL of these samples were incubated for 90 min at 37 °C and 900 rpm. Afterward, acetonitrile/methanol (400 µL, 1:1) was added, and the samples were cooled to 0 °C for 10 min. 600 µL buffer was added and the solution was centrifuged for 15 min at 16000 rpm.

The supernatant was centrifuged again for 15 min at 16000 rpm and 4 °C. The obtained solution was diluted with H_2_O/ACN/MeOH (1:1:1) to a suitable concentration for quantification. Shortly before injection into HPLC, the solution was centrifuged again for 2 min at 16000 rpm and 4 °C. The amount of remaining compound in the solution was quantified by ion count in SIM mode via HPLC-MS (ACN/water). The injection volume was 50 µL. Every experiment was performed in triplicate.

#### Human serum albumin (HSA) binding

Plasma protein binding was determined following the procedure of Börgel *et al.*(*53*).

#### *In vitro* stability in mouse and human serum

The serum stability of the radioligands [^18^F]MRS-4648 or [^18^F]PSB-19427 was evaluated by incubation in mouse serum at 37°C for up to 90 min. An aliquot of formulated [^18^F]MRS-4648 or [^18^F]PSB-19427 solution (20 μL) was added to a sample of mouse serum (200 μL), and the mixture was incubated at 37°C. Samples of 20 μL each were drawn after periods of 10, 30, 60 and 90 min and quenched in ice-cold acetonitrile (100 µL, DNA-grade) followed by centrifugation (3000 rpm) for ≥5 min. The supernatant was analyzed by analytical radio-HPLC B (method B). Serum stability investigations in human serum were performed analogously. HPLC traces are displayed in Fig. S1).

#### Enzyme inhibition assay

The CD73 enzyme inhibition assay of recombinant soluble CD73, membrane-bound human CD73, soluble rat CD73 and mouse CD73 was performed as previously described(*32, 38*).

#### Tissue collection and preparation

Tumors were obtained from two treatment-naïve breast cancer patients undergoing mastectomy surgery with axillary lymph node dissection at the Department of Plastic and General Surgery at Turku University Hospital (Turku, Finland). The first patient (patient “**X**”) was 54-year-old female with infiltrating ductal carcinoma (grade II, hormone receptor positive (ER 95%, PR 30%), Her2 negative, Ki-67 15%, lymph node status 6/55). The second patient (patient “**Y**”) was 44-year-old female with infiltrating ductal carcinoma with micropapillary differentiation (grade III, hormone receptor-negative, Her2 positive, Ki-67 25%, lymph node status 4/13). The collection of the tissues is part of an ongoing sample collection performed under the license ETMK 132/2016 with written consent from the patients. The excised primary tumors and metastatic lymph nodes were embedded in the cryo-mold with Tissue-Tek O.C.T. compound (Sakura Finetek Europe BV, The Netherlands), cut at 6 μm onto superfrost glass slides using a cryostat, and stored at -80°C.

#### Immunofluorescence staining

Tumor cryosections were processed for the immunofluorescence analysis of CD73 expression, as described elsewhere(*36*). Briefly, the slides were incubated overnight at +4°C in Shandon Sequenza Staining System (Thermo Scientific) with rabbit anti-human CD73 antibody (h5NT-1_L_, http://ectonucleotidases-ab.com/) diluted at 1:400 in 200 μl PBS containing 2% bovine serum albumin (BSA) and 0.1% (vol/vol) Triton X-100 (blocking buffer). The samples were washed and subsequently incubated for two hours at RT with Alexa Fluor®-633-conjugated goat anti-rabbit antibody (ThermoFisher Life Technologies), diluted in blocking buffer at 1:500. Alexa Fluor® 488-conjugated pan-cytokeratin (catalogue # MA5-18156, ThermoFisher) and Cy3-conjugated anti-smooth muscle cell-α (SMA-α, clone 1A4, Sigma-Aldrich) monoclonal antibodies were added during the incubation with secondary antibody for labeling the epithelial tumor cells and cancer-associated fibroblasts, respectively. The slides were mounted with ProLong® Gold Antifade reagent with DAPI (ThermoFisher) and examined using Zeiss LSM880 confocal microscope with Plan-Apochromat 10×/0.45 objective (Carl Zeiss GmbH, Jena, Germany).

#### *In situ* enzyme histochemistry

For the localization of AMPase activities in human tumors, the lead nitrate-based enzyme histochemistry was employed(*24, 33*). In brief, tissue cryosections were pre-incubated for 60 min at room temperature in 40 mM Trizma-maleate buffer (TMB, pH 7.3) supplemented with 250 mmol/L sucrose, the alkaline phosphatase inhibitor levamisole (2 mM) and different concentrations of CD73 inhibitors. The enzymatic reaction was then performed for 45 min at 37°C in a final volume of 20 mL of TMB containing 250 mM sucrose, 2 mM levamisole, 1.5 mM Pb(NO_3_)_2_, 1 mM CaCl_2_, 400 μM AMP, and tested CD73 inhibitors at the same concentrations. The lead orthophosphate precipitated in the course of nucleotidase activity was visualized as a brown deposit by incubating sections in 0.5% (NH4)_2_S for 10 s, followed by three washes in TMB for 5 min each. Slides were mounted with Aquatex medium (Merck, Germany). Tissue sections were also stained with hematoxylin and eosin (H&E). Whole slide imaging was performed using Pannoramic P250 Flash slide scanner (3DHistech Ltd., Budapest, Hungary) with a 20× objective. AMPase activity was determined by measuring AMP-specific brown staining intensities from the images using QuPath v.0.3.0 software(*54*). Simple tissue detection plugin was used to select whole tissue areas. Default values for hamatoxylin and DAB were used in colour deconvolution (color separation), and DAB channel was used as a proxy for AMPase activity. The average DAB intensity levels were measured as described earlier(*26*). The scripts for tissue detection, color deconvolution, and intensity analysis are shown in Supplementary Table 2.

#### Autoradiography

For the assessment of ^18^F-labeled CD73 tracer binding to human tissues, cryosections of breast tumor samples as well as sentinel lymph node samples were investigated. Cryosections were thawed and pre-incubated for 15 min at room temperature in 2-[4-(2-hydroxyethyl)piperazin-1-yl]ethanesulfonic acid buffer (HEPES). The ^18^F-labeled CD73 tracer was diluted to a radioactivity concentration of 1 MBq/mL. Blocking compounds were prepared according to the molar activity of the radiolabeled tracers [^18^F]MRS-4648 and [^18^F]PSB-19427 aiming at a 1000-fold excess of the blocking agent. For each tissue sample, neighboring sections were incubated for 20 min at room temperature with a mixture of 50µl of the radiotracer and 50µl of the classical CD73 inhibitor AMPCP (25 nmol, in PBS), or JMS04 (61 nmol, in HEPES), or HEPES buffer respectively. After incubation, slides were washed three times with PBS and Aqua bidest, mounted on a slide holder for the image acquisition, and covered with protection and scintillation foils. Digital autoradiography measurement of each slide was performed for 60 minutes (Micro Imager, Biospace, France). Acquired data were corrected for radioactive decay in reference to the start of the acquisition as well as the start of the incubation.

#### Human tumor xenograft experiments

MDA-MB-231 cells were cultivated in DMEM/GlutaMAX, supplemented with 10 % fetal calf serum, 100 U/mL penicillin, and 100 µg/mL streptomycin and L-glutamine to a final concentration of 8 mmol/L. AsPC-1 cells were cultivated in RPMI, supplemented with 10 % fetal calf serum, 100 U/mL penicillin, and 100 µg/mL streptomycin and L-glutamine to a final concentration of 8 mmol/L.1 x 10^6^ MDA-MB-231 or 1x 10^6^ AsPC-1 cells were injected subcutaneously into the shoulder region of 7 to 12 weeks old NSG (NOD.Cg-*Prkdc^scid^ Il2rg^tm1Wjl^*/SzJ) mice (Charles River Laboratories). Two tumors were inoculated per animal, and growth was followed by digital caliper measurements (volume = ½ (length * width^2^)).

#### In vivo imaging

Adult C57bl/6 (biodistribution study, 19.8±1.3 g) or tumor-bearing NSG mice (22.2±2.2 g) were anesthetized by isoflurane/O_2_, and one lateral tail vein was cannulated using a 27 G needle. A respective radiotracer (∼ 390 kBq/g body weight) was injected as a bolus *via* the tail vein, and subsequent PET scanning was performed. A ∼1000-fold excess of unlabeled JMS0414 (**6** for [^18^F]MRS-4648 ([^18^F]**2**)), unlabeled PSB-19427 (**1**), or PSB-12651 (**7**) (for [^18^F]PSB-19427 ([^18^F]**1**)) was injected into a subgroup of tumor-bearing mice 10 min before radiotracer injection in the blocking studies. PET imaging studies were carried out using a submillimeter high resolution (0.7 mm full width at half-maximum) small animal scanner (32 module quadHIDAC, Oxford Positron Systems Ltd., Oxford, UK) with a uniform spatial resolution (<1 mm) over a large cylindrical field (165 mm diameter, 280 mm axial length). List-mode data were acquired for 90 min and reconstructed into dynamic time frames using an iterative reconstruction algorithm. In a subgroup of mice, measurements were complemented by late time point PET scans for 20 min 4 h after injection. Subsequently, after PET acquisitions, the scanning bed was transferred to the computed tomography (CT) scanner (Inveon, Siemens Medical Solutions, U.S.), and a CT acquisition with a spatial resolution of 80 μm was performed for each mouse. Final acquisitions were performed as contrast-enhanced CT with i.v. injection of iopromid (Ultravist®370). Reconstructed image data sets were coregistered based on extrinsic markers attached to the multimodal scanning bed, and the in-house developed image analysis software MEDgical (EIMI) was used for quantification. Three-dimensional volumes of interest (VOIs) were defined over the respective organs in CT data sets, transferred to the coregistered PET data, and analyzed quantitatively. Regional uptake was calculated as the percentage of injected dose by dividing counts per min (cpm) in the VOI by total counts in the mouse multiplied by 100 (%ID/mL). The clearance of the radiotracers was calculated based on the PET results. PET imaging studies using [^18^F]FDG were performed analogously with a few variations. List-mode data were acquired for 60-75 min after tracer injection and reconstructed using an iterative reconstruction algorithm. Additionally, the scanned mice remained alive after PET and CT scans and were used the next day for imaging experiments with [^18^F]PSB-19427 ([^18^F]**1**) for direct comparison of [^18^F]FDG and [^18^F]PSB-19427 ([^18^F]**1**).

#### Ex vivo gamma counter measurements

Following the final PET-CT acquisition, 90 min p.i. or 260 min p.i., respectively, mice were euthanized by cervical dislocation, and a necropsy was performed. Ex vivo biodistribution of radioactivity was analyzed by scintillation counting (Wizard2 gamma counter, Perkin-Elmer Life Science) and the radioactivity in respective organs was decay-corrected and calculated as %ID per gram tissue (% ID/g).

### NMR data analysis

NMR spectra were processed with MestReNova 12.0 (MestreLab Research).

### Statistical analysis

Statistical analyses of in vivo PET, ex vivo biodistribution data were performed using GraphPadPrism 7.0 or GraphPad Pris 10.2. Statistical analyses were performed using unpaired t-test as indicated in the figure legends for each dataset.

## Supplementary Materials

Materials and Methods

Schemes S1 and S2

Experimental procedures for the synthesis of compounds **10**-**12** and **14**-**17**.

Fig. S1-S5

Tables T1 and T2

## Acknowledgments

The authors thank the Joint Berlin MX-Laboratory at BESSY II, Berlin, Germany, for beam time and assistance during synchrotron data collection as well as the Helmholtz Zentrum Berlin for travel support.

The authors thank Prof. Jean Sévigny for providing antibody against human CD73. We also thank Biocenter Finland, Cell Imaging and Cytometry Core of Turku Bioscience Centre and Medisiina Imaging Core, University of Turku, for imaging instrumentation.

We thank Stefanie Bouma, Sarah Köster, Roman Priebe, Christine Bätza, and Dirk Reinhardt for their technical support and assistance in animal handling and PET imaging studies. We thank the Interdisciplinary Center for Clinical Research (IZKF, core unit PIX), Münster, for their support in PET imaging studies.

## Funding

German Research Foundation (DFG) Emmy Noether program JU 2966/2-2 (AJ) German Research Foundation (DFG) SFB 1328 (CEM)

German Research Foundation (DFG) - EXC 2180 – 390900677 (AJ)

## Author contributions

Conceptualization: AJ, CEM

Methodology: AJ, CEM

Investigation: AJ, CD, SW, LG, JD, AI, CCS, GR, MS

Visualization: SS, SH, GY

Funding acquisition: AJ, CEM

Project administration: AJ

Supervision: AJ, KAJ, CEM, SS

Writing – original draft: AJ, CEM

Writing – review & editing: AJ, CEM, with support from all coauthors, who also reviewed and edited the manuscript.

## Competing interests

AJ, KJ, CEM are coinventors on patent no. WO2020/037275A1.

AJ, CD, CEM, SS, SH, SW are coinventors on patent no. PCT/EP2023/054046.

## Data and materials availability

All data are available in the main text or the supplementary materials.

