## Supporting Information for "Positron emission tomography (PET) tracer enables imaging of high CD73 expression in cancer"

**Scheme S1: Synthesis of PSB-19427 (1) and precursor 3.**

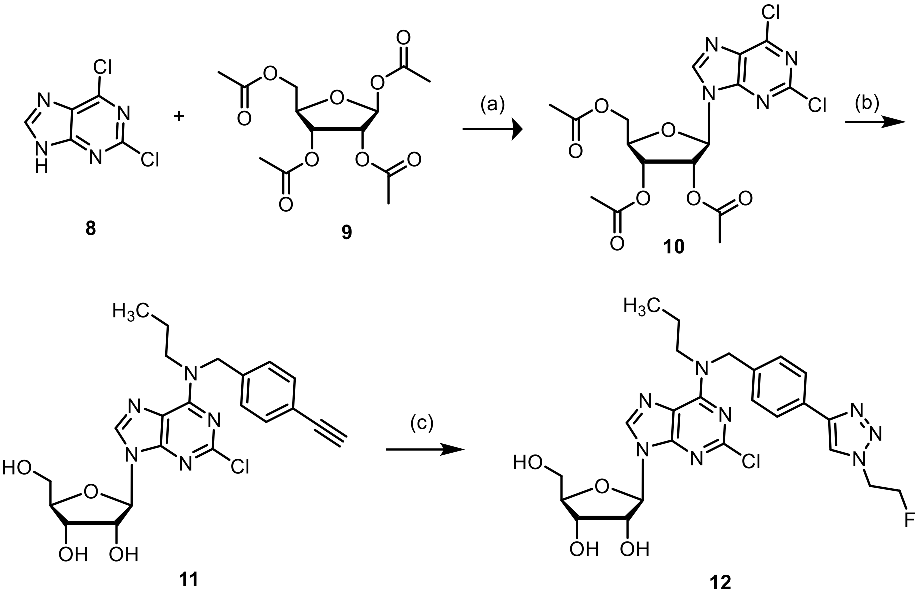

Reagents and conditions: (a) Trifluoromethanesulfonic acid, 90,000 Pa (0.9 bar), 85 °C to rt, 1 h; (b) *N*-(4-ethynylbenzyl)propan-1-amine, triethylamine, ethanol, reflux, rt, overnight; (d) 1 M $NaOCH_3$, methanol, rt, overnight; (c) 2-fluoroethyl azide, sodium ascorbate, $CuSO_4$, TBTA, DMF/$H_2O$, overnight, rt.

**Scheme S2: Synthesis of MRS-4648 (2) and precursor 4.**

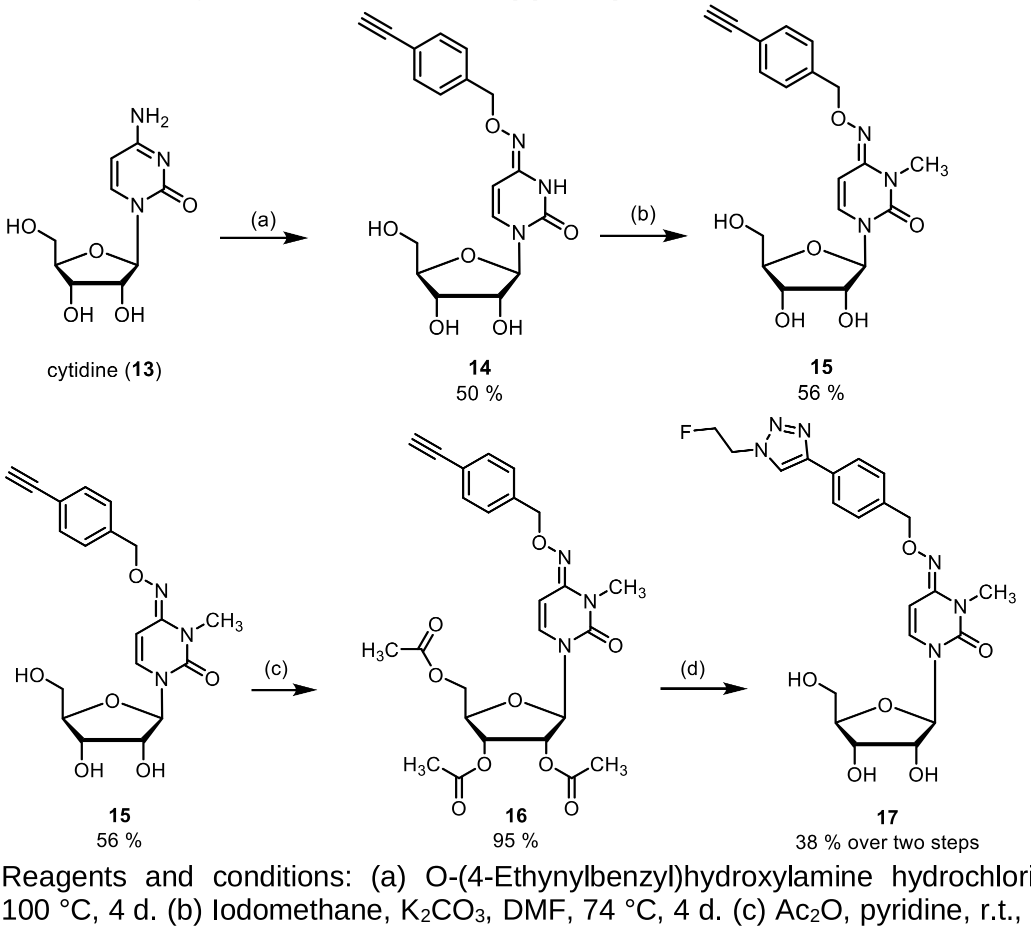

Reagents and conditions: (a) O-(4-Ethynylbenzyl)hydroxylamine hydrochloride, pyridine, 100 °C, 4 d. (b) Iodomethane, $K_2CO_3$, DMF, 74 °C, 4 d. (c) $Ac_2O$, pyridine, r.t., overnight. (d)

1) Fluoroethylazide, CuI, sodium ascorbate, $NEt_3$, DMF, r.t., overnight. 2) $NH_3$, methanol, r.t., overnight.

### Experimental section

#### Preparation of triethylammonium hydrogencarbonate (TEAC) buffer

A 1 M solution of TEAC was prepared by the following procedure: to a 1 M solution of triethylamine in deoinized water was added dry ice slowly until the pH value reached approximately 8.4-8.6.

#### 2,6-Dichloro-9-(2′,3′,5′-tri-*O*-acetyl-β-D-ribofuranosyl)-9*H*-purine (10)

1,2,3,5-Tetraacetyl-β-D-ribofuranose (5 g, 16.0 mmol, 1.0 eq) was molten at 85 °C, and 2,6-dichloropurine (3.0 g, 16.0 mmol, 1.0 eq) was added while stirring. Trifluoromethanesulfonic acid (70 µl, 0.8 mmol, 0.05 eq) was added to the reaction mixture in order to catalyze the reaction. The reaction mixture was stirred at 85 °C under reduced pressure for 30 min. Analysis by thin-layer chromatography (TLC) as performed to indicate the completion of the reaction. The mixture was then cooled to rt to allow crystallization. The solid formed was filtered off. Recrystallization from absolute ethanol yielded the desired product (4.84 g, yield 69%). $^1$H NMR (600 MHz, DMSO-$d_6$) $\delta$ 8.90 (s, 1H, 8-CH), 6.30 (d, 1H, $J$ = 4.9 Hz, 1'-CH), 5.89 (q, 1H, $J$ = 5.4 Hz, 2'-CH), 5.61 (t, 1H, $J$ = 5.6 Hz, 3'-CH), 4.43-4.28 (m, 2H, 5'-$CH_2$), 4.38 (dd, 1 H, $J$ = 3.6, 12.1 Hz, 4'-CH), 2.10 (s, 3H, $CH_3$), 2.04 (s, 3H, $CH_3$), 2.00 (s, 3H, $CH_3$). $^{13}$C NMR (126 MHz, DMSO-$d_6$) $\delta$ 170.21, 169.55, 169.41, 152.96, 151.52, 150.47, 147.04, 13.43, 86.44, 80.03, 72.55, 69.94, 62.79, 20.65, 20.52, 20.38. LC-MS (*m/z*): positive mode 447.0 $[M+H]^+$. Purity determined by HPLC-UV (254 nm)-ESI-MS: 88%. mp: 160 °C.

#### (2*R*,3*R*,4*S*,5*R*)-2-{2-Chloro-6-[(4-ethynylbenzyl)(propyl)amino]-9*H*-purin-9-yl}-5-(hydroxymethyl)tetrahydrofuran-3,4-diol (11)

2',3',5'-Tri-*O*-acetyl-2,6-dichlororibofuranosylpurine (**10**, 0.34 g, 0.77 mmol, 1.0 eq) was suspended in absolute ethanol. Triethylamine (0.2 ml, 1.54 mol, 2.0 eq) and *N*-(4-ethynylbenzyl)propan-1-amine (0.26 g, 1.54 mmol, 2.0 eq) were added to the suspension that was refluxed overnight. The progress of the reaction was monitored by TLC (DCM/methanol 9/1). After TLC indicated completion of the reaction, the solvent was evaporated. The subsequent deprotection reaction was carried out using 1 M sodium methoxide in methanol (10 ml). Purification by column chromatography (methanol/DCM 1:19) yielded the desired product as a white solid (0.25 g, 71%). $^1$H NMR (600 MHz, DMSO-$d_6$) $\delta$ 8.42 (br s, 1H, 8-CH), 7.42 (d, 2H, $J$ = 8.04 Hz, $CH_{benzene}$), 7.28 (br s, 2H, $CH_{benzene}$), 5.85 (d, $J$ = 5.83 Hz, 1H, 1'-CH), 5.54 (d, 1H, $J$ = 12.20 Hz, N-$CH_2$), 5.45 (br s, 1H, OH), 5.18 (br s, 1H, OH), 5.00 (br s, 1H, OH), 4.92 (d, 1H, $J$ = 15.05 Hz, $NCH_2$), 4.51 (br s, 1H, 2'-CH), 4.12 (s, 2H, $NCH_2$), 4.05 (q, 1H, $J$ = 5.19 Hz, 3'-CH), 3.94 (q, 1H, $J$ = 3.92 Hz, 4'-CH), 3.59 (dm, 2H, 5'-$CH_2$), 1.63 (br s, 2H, $C\underline{H_2}CH_3$), 1.05 (t, 1H, $J$ = 7.02 Hz, CCH), 0.85 (t, 3H, $CH_3$). $^{13}$C NMR (126 MHz, $CD_3OD$) $\delta$ 154.38, 152.79, 151.73, 139.27, 132.03, 127.78, 120.67, 118.46, 87.47, 85.85, 83.46, 80.83, 73.84, 70.48, 61.45, 39.72, 21.88, 18.60, 11.08. LC-MS (m/z): positive mode 458.2 $[M+H]^+$. Purity determined by HPLC-UV (254 nm)-ESI-MS: 88%. mp. 110 °C.

#### (2*R*,3*R*,4*S*,5*R*)-2-[2-Chloro-6-({4-[1-(2-fluoroethyl)-1*H*-1,2,3-triazol-4-yl]benzyl}(propyl)amino)-9*H*-purin-9-yl]-5-(hydroxymethyl)tetrahydrofuran-3,4-diol (12)

Compound **11** (0.080 g, 0.017 mmol, 1.0 eq.) was dissolved in a solution of fluoroethylazide in DMF (5 ml). TBTA (0.028 g, 0.052 mmol, 0.3 eq.) was added the reaction mixture. Copper sulfate (0.008 g, 0.052 mmol, 0.3 eq.) and sodium ascorbate (0.041 g, 0.21 mmol, 1.2 eq.) were premixed in 2 ml of water and then added to the reaction mixture, which was then stirred overnight. After TLC indicated completion of the reaction, the reaction mixture was diluted with water and extracted with ethyl acetate followed by washing with an aqueous lithium chloride solution (10%). The organic phase was dried of sodium sulfate, evaporated and then purified by normal phase column chromatography (silical gel, DCM/methanol 95/5). Appropriate fractions were pooled and the eluents were evaporated to give the desired compound as a colorless solid (yield: 76%, 0.073 g). Mp. 180 - 183 °C. $^{1}$H NMR (600 MHz, DMSO-*d6*) *δ* 8.55 (s, 1H, $CH_{triazol}$), 8.49 – 8.36 (m, 1H, 8-CH), 7.80 (d, *J* = 7.0 Hz, 2H, $CH_{benzene}$), 7.42 – 7.31 (m, 2H, $CH_{benzene}$), 5.86 (d, *J* = 5.8 Hz, 1H, 1'-CH), 5.57 (d, *J* = 20.9 Hz, 1H, $NCH_2$), 5.49 (s, 1H, OH), 5.20 (s, 1H, OH), 5.03 (s, 1H, OH), 4.95 (d, *J* = 21.2 Hz, 1H, N-$CH_2$), 4.89 (t, *J* = 4.7 Hz, 1H, F-$CH_2$-C$\underline{H_2}$), 4.81 (t, *J* = 4.7 Hz, 1H, F-$CH_2$-C$\underline{H_2}$), 4.76 (t, *J* = 4.7 Hz, 1H, F-C$\underline{H_2}$), 4.72 (t, *J* = 4.7 Hz, 1H, F-C$\underline{H_2}$), 4.52 (t, *J* = 5.3 Hz, 1H, 2'-CH), 4.15 – 4.11 (m, 1H, C3'-H), 3.94 (q, *J* = 3.8 Hz, 1H, C4'-H), 3.65 (d, *J* = 11.9 Hz, 1H, 5'-$CH_2$), 3.55 (d, *J* = 12.2 Hz, 1H, 5'-$CH_2$), 1.71 – 1.57 (m, 2H, C$\underline{H_2}CH_3$), 0.90 – 0.82 (m, 3H, $CH_3$). $^{13}$C NMR (151 MHz, DMSO-*d6*) *δ* 162.45, 154.49, 154.39, 152.86, 152.73, 151.73, 146.43, 137.88, 137.41, 129.83, 128.26, 128.02, 125.54, 121.88, 118.47, 118.37, 87.46, 85.85, 82.57, 81.45, 73.84, 70.48, 61.44, 50.41, 50.28, 35.92, 30.92, 21.37, 19.86, 11.36, 10.89. $^{19}$F NMR (565 MHz, DMSO-d6) *δ* -74.22. LC/ESI-MS (m/z): positive mode 547.40 $[M+H]^+$ Purity determined by HPLC-UV (254 nm)-ESI-MS: 98%. mp. 180 – 183 °C.

**1-[(2*R*,3*R*,4*S*,5*R*)-3,4-Dihydroxy-5-(hydroxymethyl)tetrahydrofuran-2-yl]-4-{[(4-ethynylbenzyl)oxy]imino}-3,4-dihydropyrimidin-2(1*H*)-one (14)**

O-(4-Ethynylbenzyl)hydroxylamine hydrochloride (214 mg, 1.28 mmol, 1.0 eq.) and cytidine (621 mg, 2.55 mmol, 2.0 eq.) were suspended in pyridine, and the mixture was stirred at 100 °C for 4 d. The solvent was removed *in vacuo,* and the remaining pyridine was co-evaporated with water (50 mL), toluene (50 ml), and ethanol (50 mL). The residue was purified by column chromatography using 5 % methanol in $CH_2Cl_2$ to 10 % methanol in $CH_2Cl_2$ to obtain the product as pale yellow solid (237 mg, 0.63 mmol, 50 %). $R_f$ (10 % methanol in $CH_2Cl_2$): 0.33. Purity (HPLC-A): 96 % ($R_t$ = 6.9 min). mp (140 °C→200 °C, 1 °C/min): decomposition over 163 °C. HRMS (APCI): m/z = calcd. for $C_{18}H_{20}N_3O_6$ 374.1347, found 374.1329 $[M+H]^+$.

$^{1}$H NMR ($CD_3OD$, 600 MHz): δ (ppm) = 7.45-7.42 (m, 2H, 3-C*H*$_{phenyl}$ and 5-C*H*$_{phenyl}$), 7.36-7.34 (m, 2H, 2-C*H*$_{phenyl}$ and 6-C*H*$_{phenyl}$), 7.21 (d, *J* = 8.2 Hz, 1H, 6-C*H*$_{pyrimidine}$), 5.85 (d, *J* = 5.5 Hz, 1H, 1'-C*H*), 5.55 (d, *J* = 8.2 Hz, 1H, 5-C*H*$_{pyrimidine}$), 5.02 (s, 2H, C*H*$_{2\text{-benzyl}}$), 4.14 (t, *J* = 5.5 Hz, 1H, 2'-C*H*), 4.10 (dd, *J* = 5.5, 3.9 Hz, 1H, 3'-C*H*), 3.94 (h, *J* = 3.3 Hz, 1H, 4'-C*H*), 3.77 (dd, *J* = 12.1, 2.9 Hz, 1H, 5'-C*H*H), 3.68 (dd, *J* = 12.1, 3.4 Hz, 1H, 5'-CH*H*), 3.46 (s, 1H, C*H*$_{ethynyl}$).$^{13}$C NMR ($CD_3OD$, 151 MHz): δ (ppm) = 151.5 (*C*=O$_{pyrimidine}$), 146.8 (*C*=N$_{pyrimidine}$), 140.4 (*C*-1$_{phenyl}$), 133.1 (*C*-6$_{pyrimidine}$), 133.0 (2C, *C*-3$_{phenyl}$ and *C*-5$_{phenyl}$), 128.9 (2C, *C*-2$_{phenyl}$ and *C*-6$_{phenyl}$), 123.1 (*C*-4$_{phenyl}$), 98.6 (*C*-5$_{pyrimidine}$), 89.8 (*C*-1'), 86.2 (*C*-4'), 84.3 (*C*q$_{ethynyl}$), 78.7 (*C*H$_{ethynyl}$), 76.0 (*C*H$_{2\text{-benzyl}}$), 74.8 (*C*-2'), 71.7 (*C*-3'), 62.7 (*C*-5'). FTIR (neat): ṽ ($cm^{-1}$) = 3356 (m, C-H, alkynyl), 3278 (bm, O-H and N-H), 3098 (w, C-H, aromatic), 2924, 2874 (w, C-H, aliphatic), 1682, 1663 (s, C=O and C=N), 1605 (m, C=C, pyrimidine), 1504 (w, C=C, aromatic).

**1-[(2*R*,3*R*,4*S*,5*R*)-3,4-Dihydroxy-5-(hydroxymethyl)tetrahydrofuran-2-yl]-4-{[(4-ethynylbenzyl)oxy]imino}-3-methyl-3,4-dihydropyrimidin-2(1*H*)-one (15)**

Compound **14** (117 mg, 0.31 mmol, 1.0 eq.), iodomethane (39 µL, 89 mg, 0.63 mmol, 2.0 eq.) and $K_2CO_3$ (87 mg, 0.63 mmol, 2.0 eq.) were suspended in DMF and mixture was stirred at 74 °C for 4 d. The mixture was filtrated, the residue was washed with acetone (40 mL), and the filtrate's solvent was removed *in vacuo*. The residue was purified by column chromatography using 7 % methanol in $CH_2Cl_2$ to obtain the product as pale yellow solid (67 mg, 0.17 mmol, 56 %). $R_f$ (10 % methanol in $CH_2Cl_2$): 0.41. Purity (HPLC-A): 97 % ($R_t$ = 9.0 min). mp (170 °C→230 °C, 1 °C/min): 183.0 °C. HRMS (APCI): m/z = calcd. for $C_{19}H_{22}N_3O_6$ 388.1503, found 388.1496 $[M+H]^+$. FTIR (neat): ṽ ($cm^{-1}$) = 3364 (C-H, alkynyl), 3287, 3225 (m, O-H), 2967, 2862 (w, C-H, aliphatic), 2106 (w, C≡C), 1662 (s, C=O and C=N), 1577 (m, C=C, pyrimidine), 1508 (w, C=C, aromatic). Purity (HPLC-A): >99 % ($R_t$ = 9.3 min).$^{1}$H NMR ($CD_3OD$, 600 MHz): δ (ppm) = 7.44-7.42 (m, 2H, 3-C*H*$_{phenyl}$ and 5-C*H*$_{phenyl}$), 7.35-7.32 (m, 2H, 2-C*H*$_{phenyl}$ and 6-C*H*$_{phenyl}$), 7.30 (d, *J* = 8.4 Hz, 1H, 6-C*H*$_{pyrimidine}$), 6.24 (d, *J* = 8.3 Hz, 1H, 5-C*H*$_{pyrimidine}$), 5.87 (d, *J* = 5.0 Hz, 1H, 1'-C*H*), 4.99 (s, 2H, C*H*$_{2\text{-benzyl}}$), 4.14-4.12 (m, 1H, 2'-C*H*), 4.10 (dd, *J* = 5.4, 4.1 Hz, 1H, 3'-C*H*), 3.96-3.93 (m, 1H, 4'-C*H*), 3.79 (dd, *J* = 12.2, 2.9 Hz,

1H, 5'-C*H*H), 3.70 (dd, *J* = 12.2, 3.5 Hz, 1H, 5'-CH*H*), 3.46 (s, 1H, C$H_{ethynyl}$), 3.18 (s, 3H, $CH_3$). $^1$H NMR (DMSO-$d_6$, 600 MHz): δ (ppm) = 7.49-7.43 (m, 2H, 3-C$H_{phenyl}$ and 5-C$H_{phenyl}$), 7.36 (dd, *J* = 8.4, 2.9 Hz, 3H, 2-C$H_{phenyl}$ and 6-C$H_{phenyl}$ and 6-C$H_{pyrimidine}$), 6.14 (d, *J* = 8.3 Hz, 1H, 5-C$H_{pyrimidine}$), 5.78 (d, *J* = 5.7 Hz, 1H, 1'-C*H*), 5.28 (d, *J* = 5.8 Hz, 1H, 2'-O*H*), 5.04 (d, *J* = 5.0 Hz, 1H, 3'-O*H*), 5.01 (t, *J* = 5.2 Hz, 1H, 5'-O*H*), 4.98 (s, 2H, C$H_{2\text{-benzyl}}$), 4.16 (s, 1H, C$H_{ethynyl}$), 3.97 (q, *J* = 5.6 Hz, 1H, 2'-C*H*), 3.93 (td, *J* = 5.1, 3.7 Hz, 1H, 3'-C*H*), 3.80 (q, *J* = 3.5 Hz, 1H, 4'-C*H*), 3.58 (ddd, *J* = 11.9, 5.3, 3.4 Hz, 1H, 5'-C*H*H), 3.52 (ddd, *J* = 11.9, 5.1, 3.5 Hz, 1H, 5'-CH*H*), 3.07 (s, 3H, $CH_3$).$^{13}$C NMR ($CD_3OD$, 151 MHz): δ (ppm) = 151.9 (*C*=$O_{pyrimidine}$), 151.5 (*C*=$N_{pyrimidine}$), 140.6 (*C*-$1_{phenyl}$), 133.3 (*C*-$6_{pyrimidine}$), 132.9 (2C, *C*-$3_{phenyl}$ and *C*-$5_{phenyl}$), 129.1 (2C, *C*-$2_{phenyl}$ and *C*-$6_{phenyl}$), 123.0 (*C*-$4_{phenyl}$), 93.8 (*C*-$5_{pyrimidine}$), 90.8 (*C*-1'), 86.1 (*C*-4'), 84.3 (*C*$q_{ethynyl}$), 78.6 (*C*$H_{ethynyl}$), 76.3 (*C*$H_{2\text{-benzyl}}$), 75.1 (*C*-2'), 71.5 (*C*-3'), 62.6 (*C*-5'), 29.3 ($CH_3$) $^{13}$C NMR (DMSO-$d_6$, 151 MHz): δ (ppm) = 149.8 (*C*=$N_{pyrimidine}$), 149.6 (*C*=$O_{pyrimidine}$), 139.3 (*C*-$1_{phenyl}$), 132.6 (*C*-$6_{pyrimidine}$), 131.6 (2C, *C*-$3_{phenyl}$ and *C*-$5_{phenyl}$), 128.0 (2C, *C*-$2_{phenyl}$ and *C*-$6_{phenyl}$), 120.8 (*C*-$4_{phenyl}$), 91.9 (*C*-$5_{pyrimidine}$), 88.1 (*C*-1'), 84.6 (*C*-4'), 83.4 (*C*$q_{ethynyl}$), 80.8 (*C*$H_{ethynyl}$), 74.4 (*C*$H_{2\text{-benzyl}}$), 73.1 (*C*-2'), 70.0 (*C*-3'), 61.1 (*C*-5'), 28.6 ($CH_3$).

**(2*R*,3*R*,4*R*,5*R*)-2-(Acetoxymethyl)-5-(4-{[(4-ethynylbenzyl)oxy]imino}-3-methyl-2-oxo-3,4-dihydropyrimidin-1(2*H*)-yl)tetrahydrofuran-3,4-diyl diacetate (16)**

Compound **15** (481 mg, 1.24 mmol, 1.0 eq.) was dissolved in dry pyridine (10 mL), and acetic acid anhydride (0.41 mL, 444 mg, 4.35 mmol, 3.5 eq.) was added at room temperature. The solution was stirred overnight, methanol (1 mL) was added, and the solution was stirred at room temperature for 1 h. The solvent was removed *in vacuo* and co-evaporated first with toluene (10 mL) and then ethanol (10 mL). The residue was dissolved in $CH_2Cl_2$ (50 mL), and the organic layer was washed with water (2x25 mL) and brine (25 mL). The organic layer was dried over $Na_2SO_4$. Residue was purified by column chromatography using 0 % → 1 % methanol in $CH_2Cl_2$ to obtain the product as pale yellow, sticky solid (602 mg, 1.17 mmol, 95 %). Purity (HPLC-A): 90 % ($R_t$ = 16.4 min). $R_f$ (1 % methanol in $CH_2Cl_2$): 0.45. HRMS (ESI): m/z = calcd. for $C_{25}H_{28}N_3O_9$ 514.1820, found 514.1818 $[M+H]^+$. $^1$H NMR ($CD_3OD$, 600 MHz): δ (ppm) = 7.44-7.42 (m, 2H, 3-C$H_{phenyl}$ and 5-C$H_{phenyl}$), 7.35-7.33 (m, 2H, 2-C$H_{phenyl}$ and 6-C$H_{phenyl}$), 7.02 (d, *J* = 8.4 Hz, 1H, 6-C$H_{pyrimidine}$), 6.29 (d, *J* = 8.3 Hz, 1H, 5-C$H_{pyrimidine}$), 5.89 (d, *J* = 5.2 Hz, 1H, 1'-C*H*), 5.41 (dd, *J* = 6.1, 5.1 Hz, 1H, 2'-C*H*), 5.37 (dd, *J* = 6.1, 5.0 Hz, 1H, 3'-C*H*), 5.00 (s, 2H, C$H_{2\text{-benzyl}}$), 4.32 (dd, *J* = 3.9, 1.7 Hz, 2H, 5'-C$H_2$), 4.28 (dt, *J* = 5.0, 3.9 Hz, 1H, 4'-C*H*), 3.46 (s, 1H, C$H_{ethynyl}$), 3.17 (s, 3H, C$H_{3\ pyrimidine}$), 2.09(8) (s, 3H, 3'-C$H_{3\ acetate}$), 2.09(5) (s, 3H, 5'-C$H_{3\ acetate}$), 2.06 (s, 3H, 2'-C$H_{3\ acetate}$). $^{13}$C NMR ($CD_3OD$, 151 MHz): δ (ppm) = 172.2 (*C*=O-5'$_{acetate}$), 171.4 (2C, *C*=O-2'$_{acetate}$ and *C*=O-3'$_{acetate}$), 151.4 (*C*=$O_{pyrimidine}$), 151.0 (*C*=$N_{pyrimidine}$), 140.5 (*C*-$1_{phenyl}$), 133.2 (*C*-$6_{pyrimidine}$), 132.9 (2C, *C*-$3_{phenyl}$ and *C*-$5_{phenyl}$), 129.1 (2C, *C*-$2_{phenyl}$ and *C*-$6_{phenyl}$), 123.0 (*C*-$4_{phenyl}$), 94.5 (*C*-$5_{pyrimidine}$), 90.3 (*C*-1'), 84.3 (*C*$q_{ethynyl}$), 80.8 (*C*-4'), 78.6 (*C*$H_{ethynyl}$), 76.4 (*C*$H_{2\text{-benzyl}}$), 73.8 (*C*-2'), 71.7 (*C*-3'), 64.4 (*C*-5'), 29.3 (*C*$H_{3\ pyrimidine}$), 20.7 and 20.4 (2C, *C*$H_3$-3'$_{acetate}$ and *C*$H_3$-5'$_{acetate}$), 20.3 (*C*$H_3$-2'$_{acetate}$). FTIR (neat): ṽ ($cm^{-1}$) = 3271 (w, C-H, alkynyl), 2932, 2870 (w, C-H, aliphatic), 1744 (s, C=O, acetate), 1686, 1662 (s, C=O and C=N, pyrimidine), 1585 (m, C=C, pyrimidine), 1504 (w, C=C, aromatic).

**1-[(2*R*,3*R*,4*S*,5*R*)-3,4-Dihydroxy-5-(hydroxymethyl)tetrahydrofuran-2-yl]-4-[({4-[1-(2-fluoroethyl)-1*H*-1,2,3-triazol-4-yl]benzyl}oxy)imino]-3-methyl-3,4-dihydropyrimidin-2(1*H*)-one (17)**

2-Fluroethanol (191 µL, 210 mg, 3.28 mmol, 3.72 eq.), 4-toluenesulfonyl chloride (687 mg, 3.61 mmol, 4.10 eq.) and KOH (276 mg, 4.92 mmol, 5.59 eq.) were suspended in dry $CH_2Cl_2$ (5 mL), and the mixture was stirred at room temperature for 3 h. $CH_2Cl_2$ (22 mL) was added, and the mixture was extracted with water (2x25 mL). The organic layer was dried over $Na_2SO_4$, and the solvent was removed, yielding crude 2-fluoroethyl 4-methylbenzenesulfonate as a pale yellow liquid (734 mg, 3.36 mmol, 103 %), which was used without further purification

$^1$H NMR ($CDCl_3$, 400 MHz, 367M): δ (ppm) = 7.85-7.76 (m, 2-C$H_{tosyl}$ and 6-C$H_{tosyl}$), 7.40-7.31 (m, 2H, 3-C$H_{tosyl}$ and 5-C$H_{tosyl}$), 4.66-4.61 (m, 1H, FC*H*HC$H_2$O), 4.53-4.49 (m, 1H,

FCH*H*CH$_2$O), 4.33-4.28 (m, 1H, FCH$_2$C*H*HO), 4.25-4.21 (m, 1H, FCH$_2$CH*H*O), 2.46 (s, *3*H, C*H*$_3$). $^{13}$C NMR (CDCl$_3$, 101 MHz, 367M): δ (ppm) = 145.3 (*C*-4$_{tosyl}$), 132.8 (*C*-1$_{tosyl}$), 130.1 (2C, *C*-3$_{tosyl}$ and *C*-5$_{tosyl}$), 128.1 (2C, *C*-2$_{tosyl}$ and *C*-6$_{tosyl}$), 80.7 (d, *J* = 173.6 Hz, F*C*H$_2$CH$_2$O), 68.6 (d, *J* = 20.9 Hz, FCH$_2$*C*H$_2$O), 21.8 (CH$_3$). $^{19}$F NMR (CDCl$_3$, 376 MHz, 367M): δ (ppm) = -224.7 (tt, *J* = 47.1, 27.1 Hz).

2-Fluoroethyl 4-methylbenzenesulfonate was dissolved in dry DMF (50 mL), and $NaN_3$ (1.28 g, 19.67 mmol, 22.35 eq.) was added. The mixture was stirred at 60 °C for 2 d and was then filtrated. CuI (624 mg, 3.28 mmol, 4.10 eq.), sodium ascorbate (650 mg, 3.28 mmol, 4.10 eq.) and triethylamine (0.45 mL, 332 mg, 3.28 mmol, 4.10 eq.), followed by compound **16** (451 mg, 0.88 mmol, 1.0 eq.) in DMF (5 mL) were added to the filtrate. The mixture was stirred at room temperature overnight, controlled by LC/MS. The solvent was removed in vacuo, and the residue was suspended in ethyl acetate (100 mL). The organic layer was washed with aqueous sodium tartrate (1 M, 2x50 mL) and water (50 mL). Organic layer was dried over $Na_2SO_4$, and the solvent was removed *in vacuo*, yielding crude (2*R*,3*R*,4*R*,5*R*)-2-(acetoxymethyl)-5-{4-[({4-[1-(2-fluoroethyl)-1*H*-1,2,3-triazol-4-yl]benzyl}oxy)imino]-3-methyl-2-oxo-3,4-dihydropyrimidin-1(2*H*)-yl}tetrahydrofuran-3,4-diyl diacetate (516 mg, 0.86 mmol, 97 %), which was used without further purification. The intermediate was dissolved in methanol, and $NH_3$ (4 M in methanol, 3.3 mL, 13.2 mmol, 15 eq.) was added. The mixture was stirred overnight, controlled by LC/MS. The solvent was removed *in vacuo*, and the residue was purified *via* semi-preparative C18-HPLC (RP-HPLC-C), yielding the product (132 mg, 0.34 mmol, 39 %) as a pale yellow solid. $R_t$ (RP-HPLC-C): 30.5-32.5 min: solvent A: $CH_3CN$, solvent B: water, 0-2 min: 15 % solvent A, 3 mL/min to 5 mL/min, 2-50 min: 15 % to 100 % solvent A, 50-55 min: 100 % solvent A, 55-56 min: 100 % to 15 % solvent A, 56-60 min: 15 % solvent A, 60-61 min: 15 % solvent A, 5 mL/min to 0.1 mL/min, Injection volume: 800 µL. Purity (HPLC-A, 254 nm): 96 % ($R_t$ = 7.4 min). HRMS (ESI): m/z = calcd. for $C_{21}H_{26}FN_6O_6$ 477.1892, found 477.1930 $[M+H]^+$. $^1$H NMR (CD$_3$OD, 400 MHz): δ (ppm) = 8.35 (s, 1H, C*H*$_{triazole}$), 7.98 (s, 2H, C*H*$_{DMF}$), 7.83-7.79 (m, 2H, 3-C*H*$_{phenyl}$ and 5-C*H*$_{phenyl}$), 7.47-7.43 (m, 2H, 2-C*H*$_{phenyl}$ and 6-C*H*$_{phenyl}$), 7.29 (d, *J* = 8.4 Hz, 1H, 6-C*H*$_{pyrimidine}$), 6.26 (d, *J* = 8.3 Hz, 1H, 5-C*H*$_{pyrimidine}$), 5.87 (d, *J* = 4.9 Hz, 1H, 1'-C*H*), 5.02 (s, 2H, C*H*$_{2\text{-}benzyl}$), 4.92 (dd, *J* = 5.3, 4.1 Hz, 1H, CH$_2$C*H*HF), 4.82-4.78 (m, 2H, CH$_2$CH*H*F and C*H*HCH$_2$F), 4.76-4.73 (m, 1H, CH*H*CH$_2$F), 4.15-4.09 (m, 2H, 2'-C*H* and 3'-C*H*), 3.94 (q, *J* = 3.4 Hz, 1H, 4'-C*H*), 3.79 (dd, *J* = 12.2, 2.9 Hz, 1H, 4'-C*H*H), 3.69 (dd, *J* = 12.1, 3.5 Hz, 1H, 4'-CH*H*), 3.19 (s, 3H, C*H*$_{3\ pyrimidine}$), 2.99 (s, 6H, C*H*$_{3\ DMF}$), 2.86 (d, *J* = 0.7 Hz, 3H, C*H*$_{3\ DMF}$). $^{13}$C NMR (CD$_3$OD, 101 MHz): δ (ppm) = 164.9 (*C*=O$_{DMF}$), 151.9 (*C*=O$_{pyrimidine}$), 151.5 (*C*=N$_{pyrimidine}$), 148.9 (*C*-1$_{triazole}$), 140.0 (*C*-1$_{phenyl}$), 133.2 (*C*-6$_{pyrimidine}$), 131.0 (*C*-4$_{phenyl}$), 129.9 (2C, *C*-3$_{phenyl}$ and *C*-5$_{phenyl}$), 126.6 (2C, *C*-2$_{phenyl}$ and *C*-6$_{phenyl}$), 122.9 (*C*-2$_{triazole}$), 93.9 (*C*-5$_{pyrimidine}$), 90.8 (*C*-1'), 86.1 (*C*-4'), 82.8 (d, *J* = 170.8 Hz, *C*H$_2$F), 76.5 (*C*H$_{2\text{-}benzyl}$), 75.0 (*C*-2'), 71.5 (*C*-3'), 62.7 (*C*-5'), 52.0 (d, *J* = 20.3 Hz, *C*H$_2$CH$_2$F), 37.0 (2C, *C*H$_{3\ DMF}$), 31.7 (2C, *C*H$_{3\ DMF}$), 29.3 (CH$_{3\ pyrimidine}$). $^{19}$F NMR (CD$_3$OD, 376 MHz): δ (ppm) = -224.5 (tt, *J* = 46.8, 27.0 Hz). FTIR (neat): ṽ (cm$^{-1}$) = 3345 (bm, N-H and O-H), 3140 (w, C-H, alkene), 2963 and 2924 (w, C-H, aliphatic), 1659 (s, C=O and C=N), 1585 (m, C=C, pyrimidine), 1497 (w, C=C, aromatic).

**Fig S1: *In vitro* stability of the developed PET tracers in mouse and human serum.**

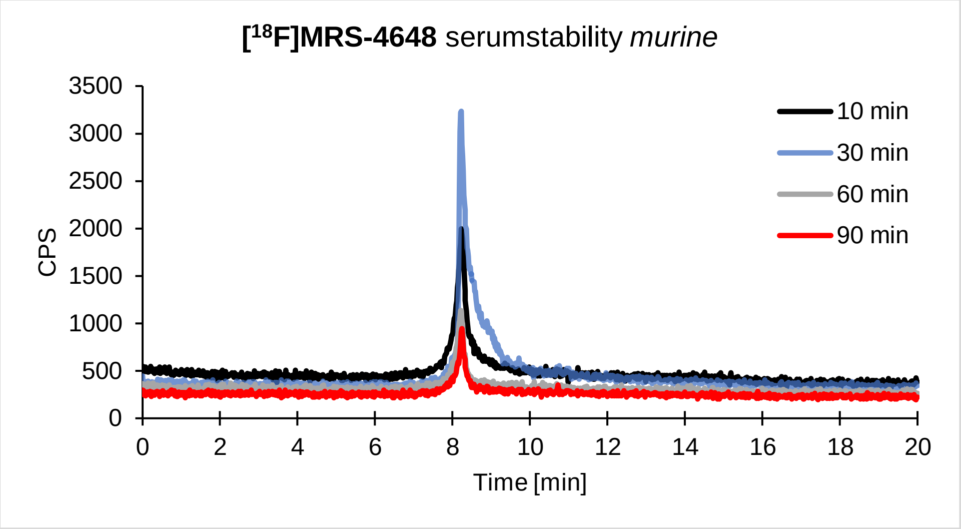

Mouse serum stability of [$^{18}$F]MRS-4648.

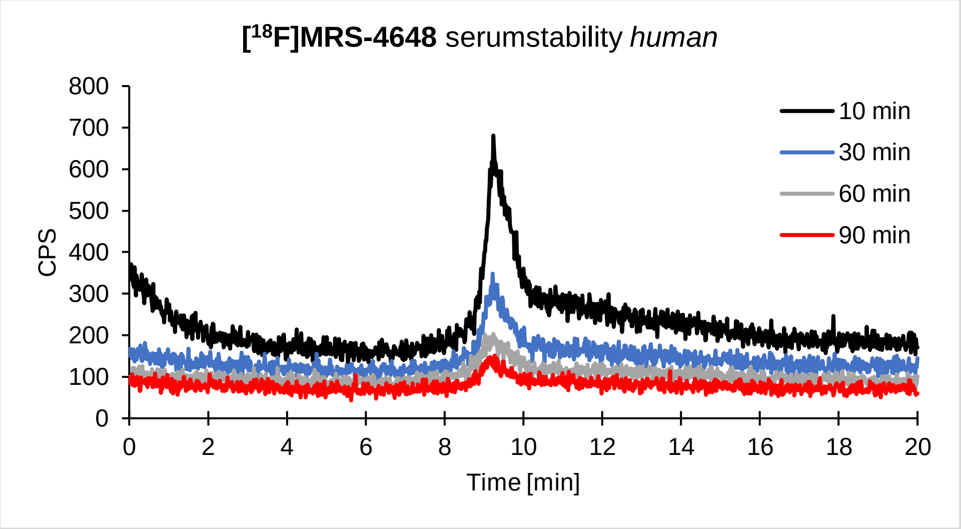

Human serum stability of [$^{18}$F]MRS-4648.

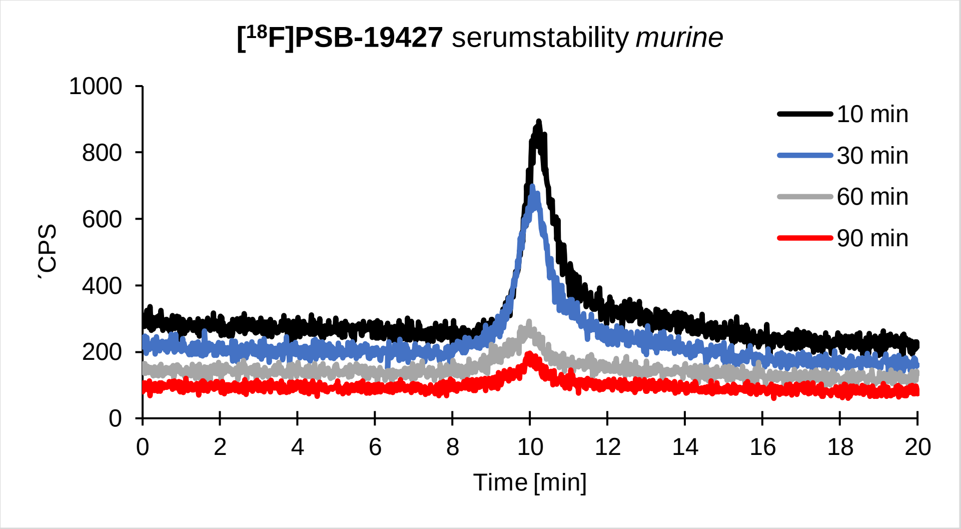

Mouse serum stability of [$^{18}$F]PSB-19427.

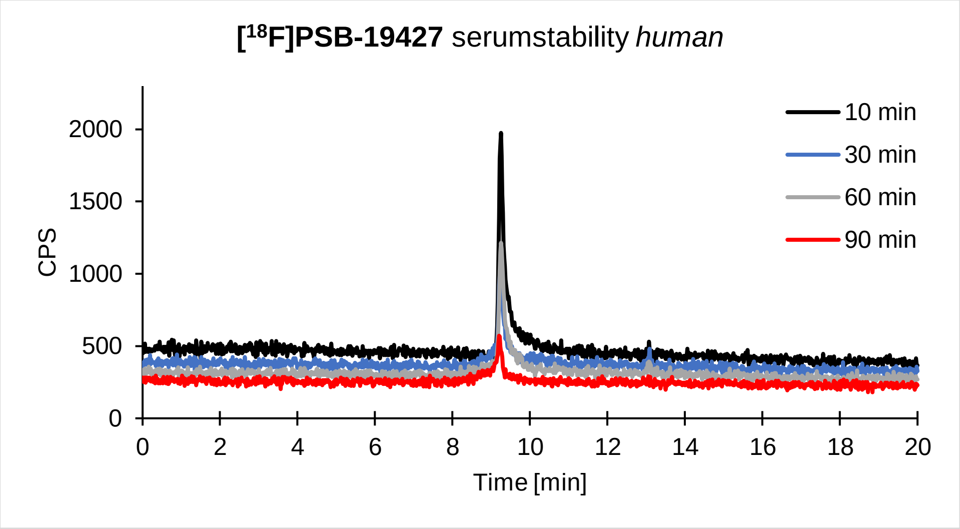

Human serum stability of [$^{18}$F]PSB-19427.

**Fig. S2: Dynamic tracer uptake of [$^{18}$F]PSB-19427 ([$^{18}$F]1) in two MDA-MB-231 tumor-bearing mice**

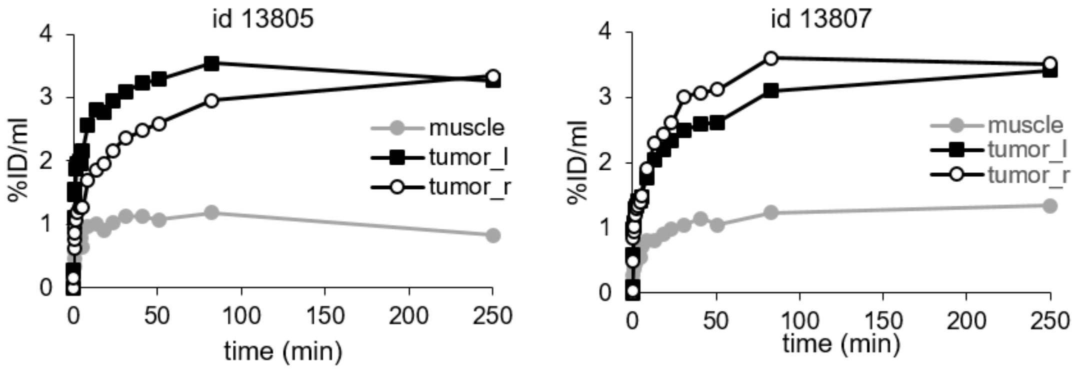

**Fig S3: Fig S2: Quantitative PET analysis of [$^{18}$F]1 uptake: concentration of [$^{18}$F]1 after 90 and 260 min expressed as tumor-to-muscle ratios.**

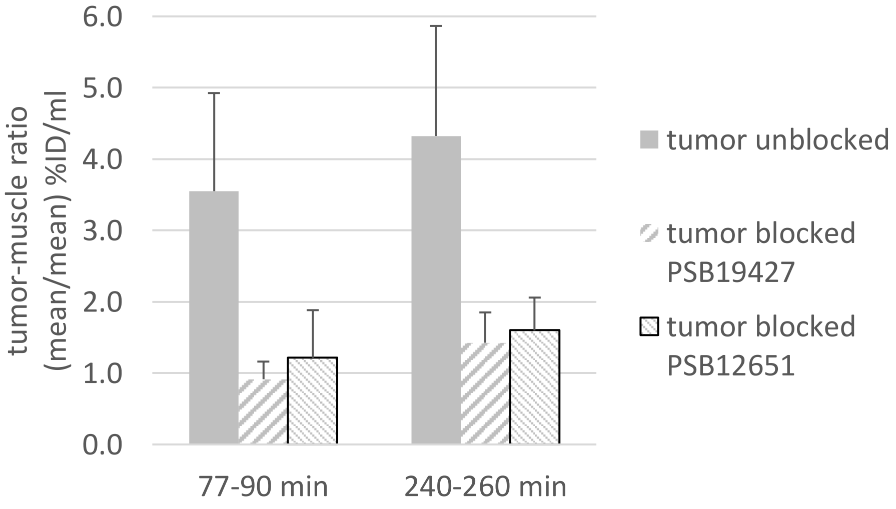

**Figure S4: Comparison of inhibitor binding modes.**

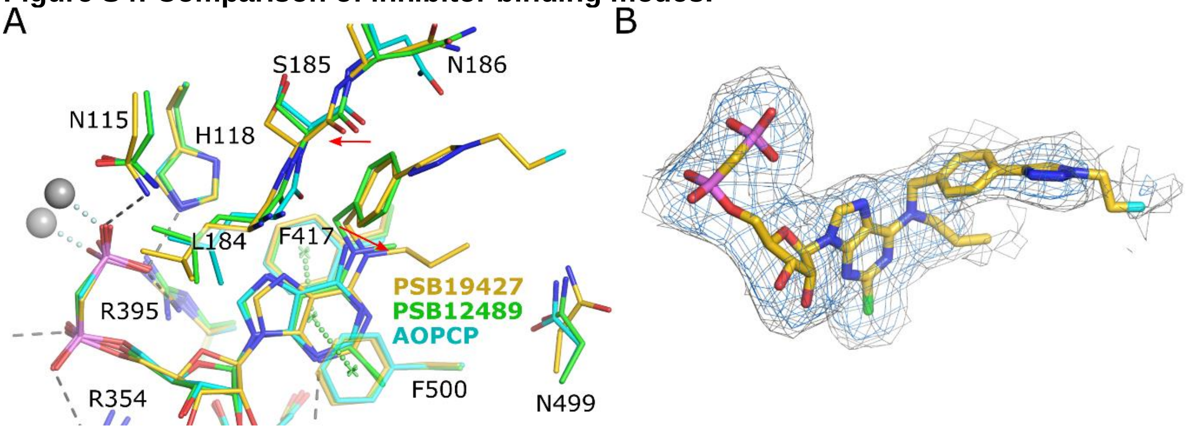

(A) Superposition of the binding modes of PSB19427, PSB12489 (pdb id 6s7h) and AOPCP (4h2i). The two red arrows mark the movements of the N6 atom of the inhibitors and of the region 184-196. The structures have been superimposed based on the Cα atoms of the C-terminal domain. (B) Polder omit electron density map of PSB12489 contoured at 4.0 σrms (blue) and at 3.0 σrms (gray).

**Table S1: Details of crystallographic data collection and refinement**

| | **9HD5 (PSB19427)** |
|---|---|
| PDB code | 9HD5 |
| **Crystallization and data collection** | |
| Crystallization buffer | 18% PEG 6000; 0.1 M BisTrisPropane pH 7.0, 10 mM PSB19427, 10 µM $ZnCl_2$ |
| Crystallization drop | 1 µL crystallization buffer + 1 µL of 7 mg/mL CD73 in 10 mM Tris pH 8.0, 10 µM $ZnCl_2$, 10 mM A830 |
| Cryobuffer | cryst. buffer + 20 % PEG200 |
| Wavelength (Å) | 0.91841 |
| Resolution range (Å)[1] | 49.37-2.91 (3.08-2.91) |
| Space group | P $2_1$ $2_1$ 2 |
| Unit cell parameters a, b, c (Å) | 92.63 233.36 54.06 |
| Unique reflections | 16993 (849) |
| Multiplicity | 12.5 (11.3) |
| Completeness (%) | 90.8 (65.4) |
| Mean I/sigma(I) | 5.5 (1.1) |
| R-meas | 0.992 (8.493) |
| R-pim | 0.279 (2.606) |
| R-merge | 0.951 (8.048) |
| CC1/2 | 0.977 (0.122) |
| Resolution aniso (Å)* | 4.38, 2.81, 2.85 |
| Wilson B-factor ($Å^2$) | 56.9 |

| | |
|---|---|
| **Refinement** | |
| Resolution range (Å) | 49.37-2.911 (2.99-2.91) |
| R-work | 0.1896 (0.2488) |
| R-free | 0.2725 (0.4649) |
| Mean B-factor ($Å^2$) | 57.7 |
| Number of non-hydrogen atoms | |
| Protein | 8005 |
| Heterogen (ligands, metals) | 96 |
| solvent | 118 |
| Root mean square deviations | |
| Bonds (Å) | 0.008 |
| Angles (°) | 1.06 |
| Ramachandran statistics (Molprobity) | |
| Favored (%) | 93.12 |
| Allowed (%) | 6.0 |
| Outliers (%) | 0.88 |

*Diffraction limits along a*, b* and c* as determined by the Staraniso algorithm

**Table S2. QuPath v.0.3.0 scripts for staining intensity measurement.**

| |
|---|
| A<br>runPlugin('qupath.imagej.detect.tissue.SimpleTissueDetection2', '{"threshold": 254, "requestedPixelSizeMicrons": 20.0, "minAreaMicrons": 2000000.0, "maxHoleAreaMicrons": 1000000.0, "darkBackground": false, "smoothImage": true, "medianCleanup": true, "dilateBoundaries": false, "smoothCoordinates": true, "excludeOnBoundary": true, "singleAnnotation": false}'); |
| B<br>setColorDeconvolutionStains('{"Name" : "H-DAB default", "Stain 1" : "Hematoxylin", "Values 1" :"0.65111 0.70119 0.29049 ", "Stain 2" : "DAB", "Values 2" : "0.26917 0.56824 0.77759 ", "Background" :" 255 255 255 "}'); |
| C<br>runPlugin('qupath.lib.algorithms.IntensityFeaturesPlugin', '{"pixelSizeMicrons": 2.0, "region": "ROI", "tileSizeMicrons": 25.0, "colorOD": false, "colorStain1": false, "colorStain2": true, "colorStain3": false, "colorRed": false, "colorGreen": false, "colorBlue": false, "colorHue": false, "colorSaturation": false, "colorBrightness": false, "doMean": true, "doStdDev": false, "doMinMax": false, "doMedian": false, "doHaralick": false, "haralickDistance": 1, "haralickBins": 32}'); |

(A) Whole tissue areas were detected with the Simple Tissue Detection function. (B) The default DAB color deconvolution values were used as a proxy for AMPase activity (brown deposit) analysis. (C) The intensities on the DAB channel were measured from the whole tissue areas using the Intensity Features function.
